# Efficient elastic tissue motions indicate general motor skill

**DOI:** 10.1101/2025.02.25.636457

**Authors:** Praneeth Namburi, Roger Pallarès-López, Duarte Folgado, Uriel Magana-Salgado, Enya Ryu, Armin Kappacher, Hugo Gamboa, Brian W. Anthony, Luca Daniel

## Abstract

Insights into the general nature of motor skill could fundamentally change how we develop movement abilities, with implications for musculoskeletal well-being and injury. Here, we sought to identify indicators of general motor skill–those shared by experts across disciplines (e.g., squash, ballet, volleyball) during non-specialized movements (e.g., reaching for water). Identifying such general indicators of motor skill has remained elusive. Using ultrasound imaging with deep learning and optical flow analysis, we tracked elastic tissues (muscles and associated connective tissues) during a simple reaching task performed similarly by world-class athletes and regional-level athletes drawn from diverse disciplines, as well as untrained non-experts. We analyzed previously unexamined inefficiencies: transverse elastic tissue motions orthogonal to muscle fiber direction and physiological tremors, which are oscillations that do not contribute to the net work done by muscles. We discovered that world-class experts minimize both these inefficient motions compared to regional level athletes and non-experts. While regional-level athletes surprisingly showed similar inefficiencies to non-experts, they used elastic tissues more effectively, achieving equivalent movements with smaller actuation-related tissue motions. We establish elastic tissue motion as a key indicator of general motor skill, expanding our understanding of elastic mechanisms and their role in general aspects of motor skill.

## Introduction

Following Spearman’s work on formalizing a measure of general intelligence in 1904 [1], there was great enthusiasm to measure general motor skill [2,3]. However, subsequent studies showed that performance in one sport (e.g., volleyball) does not correlate with performance in another (e.g., tennis) [4–10]. This has shaped the current thinking that motor skills are highly specific, shaped by the unique demands and training methods of each discipline [11].

This perspective has directed research toward identifying domain-specific indicators of expertise [12–17]. For example, expert golfers exhibit smoother golf swings [12], expert weightlifters generate greater force with less muscle activity [17], expert ballet dancers and gymnasts show reduced variability in their movements and muscle activity across task repetitions [13,14], and expert runners display lower variability in the oscillation of their center of mass [15]. Therefore, the current perspective steers away from exploring general aspects of motor skill. We contend that the absence of performance correlation across disciplines [4–10] does not preclude the existence of mechanisms contributing to general motor expertise.

Therefore, it remains unclear whether there exist fundamental mechanisms that influence movement efficiency, quality, and injury risk. We argue that if such mechanisms exist, experts across diverse disciplines (e.g., squash, ballet, volleyball) would exploit them more effectively, resulting in shared indicators of general motor skill among elite athletes across sports and performance arts. Although harnessing these mechanisms alone may not suffice to improve performance without domain-specific training, they could reveal key skill-related optimizations [18–22] made by the motor system—that is, optimizations performed by a motor system that is able to meet high demands, regardless of the specific nature of the demands. This knowledge could be applied to enhance musculoskeletal health in the general population.

Beyond generalizing across movement disciplines, we believe these fundamental mechanisms, if they exist, should be evident in non-specialized movements as well. Expert athletes undergo long and rigorous training that changes fundamental aspects of their body such as heart wall thickness [23] and muscle fiber composition [24]. Even a short amount of strength training changes how muscle fibers are recruited by the nervous system [25,26]. Such fundamental changes should logically extend beyond specialized movements into everyday actions. Yet, *it is unknown the extent to which such changes manifest in simple, everyday movements, such as reaching for a glass of water, made by trained and untrained people alike*.

In this paper, <u>we seek to identify indicators of general motor skill that unify experts across</u> <u>domains and manifest in everyday actions</u>. These shared characteristics could offer valuable insights into the physical substrates and mechanisms underlying the general nature of motor skill.

Our novel idea is that indicators of general motor skill are not evident in the external appearance of the movement – the more obvious motions of body segments – but rather in the internal *motions of tissues* producing the movement. For instance, when reaching for a glass of water, an expert’s movement may appear similar to that of an untrained individual, yet the expert’s elastic tissues may move differently when performing the task.

We hypothesize that experts’ elastic tissues move less than those of intermediates and non-experts when performing similar body segment motions. Experts are those who have achieved national or international-level recognition, intermediates are those who have achieved regional level recognition, and non-experts are those who have not systematically honed specific motor skills (see Expertise Categorization). In this paper, we use the term muscles and elastic tissues to refer to skeletal muscles and associated connective tissues.

Our innovation is to look at muscles not merely as actuators, but to consider their movements more broadly. Using a novel combination of optical flow [27] and deep learning algorithms [28,29] on ultrasound videos of moving tissues, we examine three types of muscle motions – muscle length changes, transverse motions of muscles in the plane orthogonal to the direction of the muscle fibers, and physiological tremors. Here, *<u>we show that each of these muscle motions unites experts across various domains, and</u> <u>distinguishes them from non-experts</u>*.

**<u>Conceptualization: three types of muscle motions</u>** (Fig. 1). Muscles change length along the direction of muscle fibers due to forces acting on the muscle and forces generated by the muscle. The role of ***muscle length changes*** in actuating body segments has been extensively studied in both animals and humans [30,31].

**Fig. 1.**
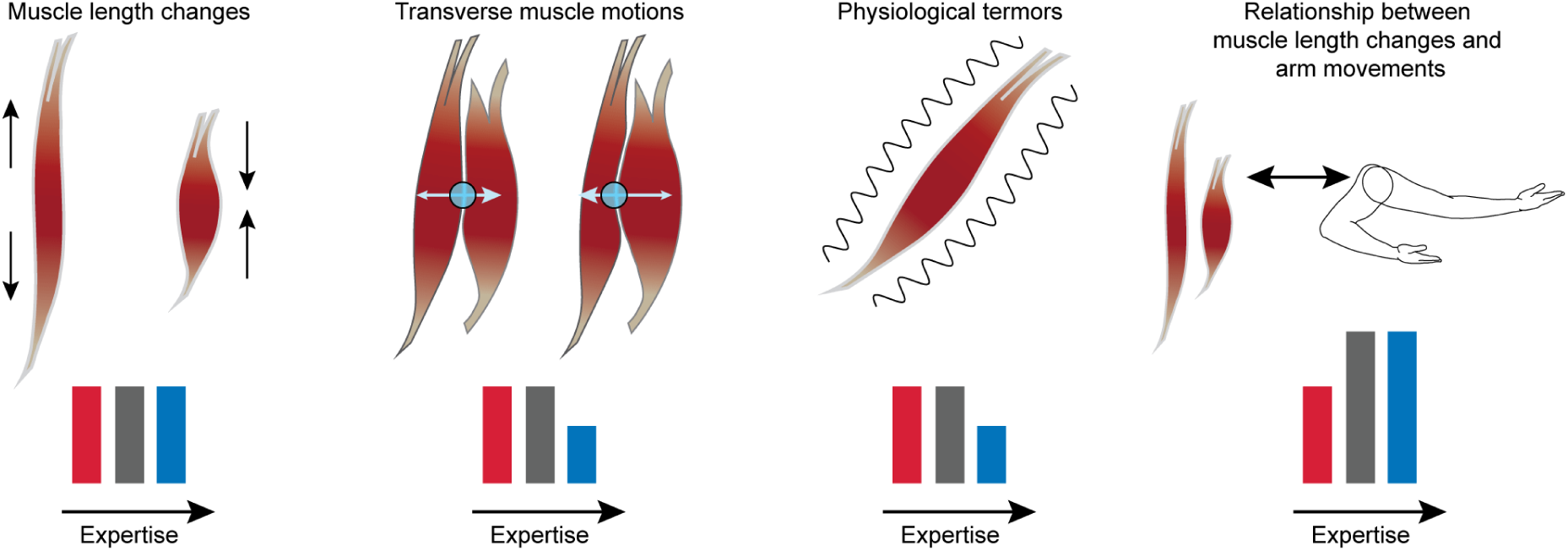
Conceptualization and a graphical summary of the paper’s main findings. The three categories of muscle motions examined in this study are as follows: 1. Muscle length changes. 2. Transverse muscle motions, which reflect adjustments due to constraints imposed by surrounding tissues such as bone, connective tissue, and fat tissue. These are measured by tracking the boundary between two muscles. 3. Physiological tremors, which arise due to neuromechanical interactions. In individuals at the highest levels of movement expertise, regardless of the movement discipline, we observed reduced transverse muscle motions and physiological tremors, as depicted in the bottom row. The red, gray, and blue bars depict findings related to the non-expert, intermediate, and expert groups, that is, participants without formal training, regional level athletes, and national/international level athletes (see Expertise Categorization in Methods). Finally, the relationship between muscle length changes and arm movements is more effective for both intermediates and experts. The takeaway from this study is that muscle motions indicate movement expertise across domains, and achieving expertise in movement impacts muscle motions beyond the specialized movements learned in any movement discipline. Muscle motions are effective at the intermediate level and more efficient (reduced transverse muscle motions and tremors) in a movement expert.

However, muscles also move in the plane orthogonal to the direction of the muscle fibers. These other movements reflect adjustments in response to the constraints imposed by surrounding structures such as bones, fat, skin, and adjacent muscles. We define these adjustments as ***transverse muscle motions*** in contrast to longitudinal muscle length changes. To our knowledge, these transverse motions have not been studied before, likely because muscles are primarily viewed as actuators [32].

During movement, healthy individuals typically experience subtle, high-frequency oscillations known as **physiological tremors** [33–36]. The 8-12 Hz component of physiological tremors is readily detected on the skin, and has a multifactorial etiology based on the mechanical properties of tissues, and the nervous system [37–39].

Physiological tremors are benign [40], often invisible to the naked eye [40], and do not compromise the motion of body segments, unlike some pathological tremors associated with conditions such as Parkinson’s disease [37,41]. In this paper, “tremor” refers only to the 8-12 Hz component of the physiological tremor [41,42] and not to any other type of tremor [40].

Transverse muscle motions occur orthogonal to the direction of force generation and therefore do not contribute to the work done by the muscle. Similarly, physiological tremors are oscillatory movements that do not contribute to the net work done by the muscle. Therefore, we denote transverse muscle motions and physiological tremors as ***inefficiencies*** in muscle motions. We refer to muscle motions as being more ***efficient*** when transverse muscle motions and tremors are minimized. Note that this is different from metabolic efficiency, which is not studied in this work. Lastly, we define muscle length changes as being more ***effective*** when the ratio of body segment motions to muscle length changes is higher – that is, when smaller muscle length changes correspond to similar body segment motions. Our findings reveal that <u>intermediates and</u> <u>experts have more effective muscle length changes than non-experts, and experts have</u> <u>more efficient muscle motions than both intermediates and non-experts.</u>

## Results

### Task design and validation

To contrast experts’ muscle motions to those of intermediates and non-experts, we used an unconstrained reaching task (Movie S1) that evokes similar arm movements across participant groups in terms of gross kinematics. This task was selected to avoid complexity and strength components that might be expected to vary among subjects in diverse movement disciplines. To understand the goal-agnostic characteristics of muscle motions, we designed this reaching task without a physical target (goal). We instructed the participants, who were standing, to reach forward in front of them, starting their hand near their hip. To ensure kinematic comparison across groups, movements were paced using a visual metronome. We set the metronome pace to a period of 6 seconds, with equal time for extension and retraction. Since humans are worse at performing slow movements [43,44], we reasoned that a slow pace might better reveal expertise-related indicators in muscle motions.

To check whether the designed reaching task evoked similar arm motions across participant groups, we compared the speed, variability, and smoothness of arm movements. Speed, obtained from five tracked points along the arm (Fig. 2a, Fig. S1) was similar across the three participant groups (Fig. 2b-c). Variability in arm speed across reach cycles, and movement smoothness were also similar across the three groups (Fig. S2), establishing that the kinematic characteristics of arm motion were similar across levels of movement expertise.

**Fig. 2.**
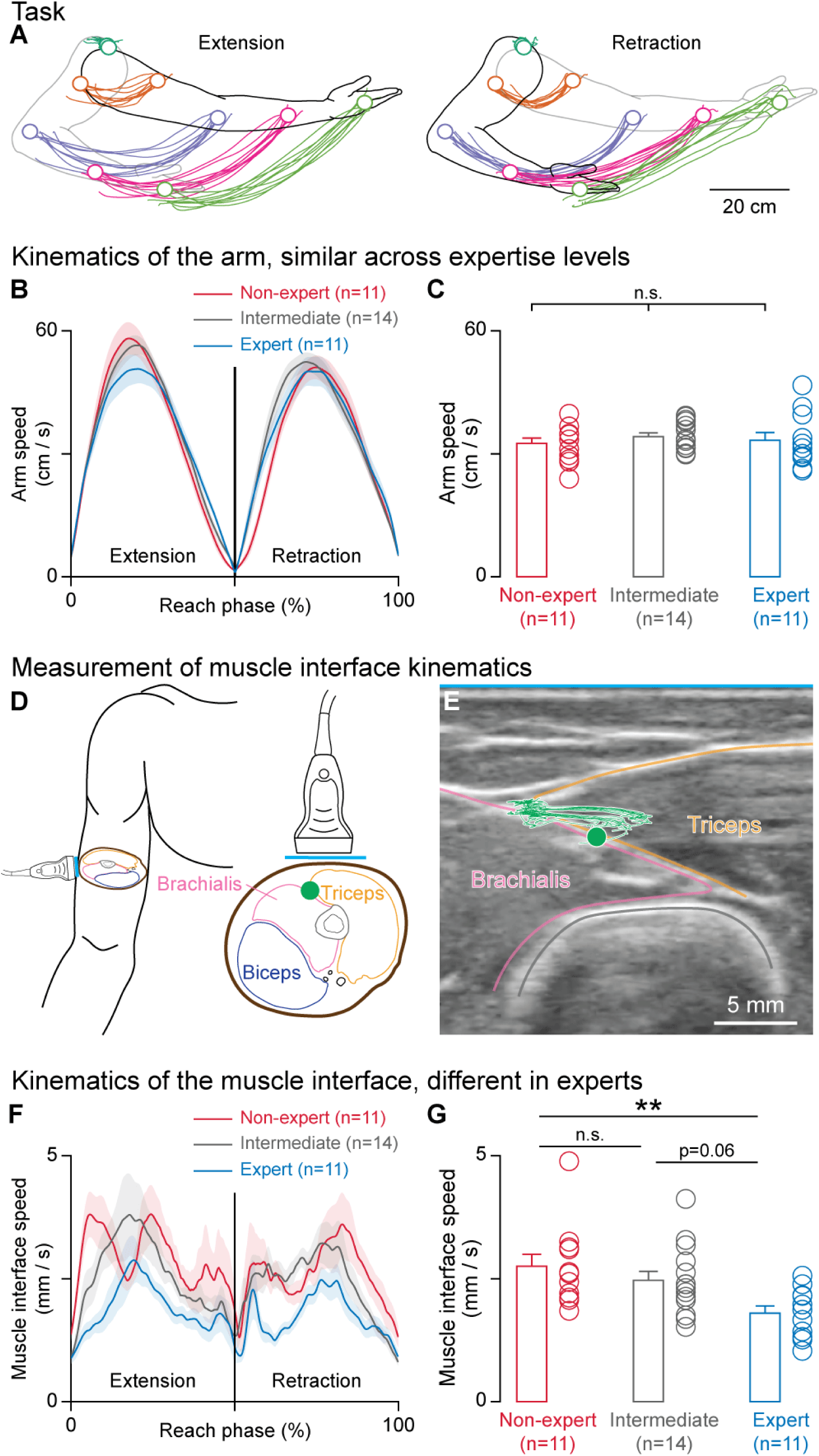
Movement experts have reduced transverse muscle motions, even when arm movements are similar to those of non-experts and intermediates. (**A**-**C**) Measurement and quantification of arm movement. (**A**) As participants performed a reaching task, the motion paths of five points along the arm were tracked using optical motion capture. The positions of these markers and example paths from one participant are shown. (**B**) Arm speed was estimated using the first derivative of the first principal component of the positional data from all five markers. The arm speed profiles from each of the participant groups, non-expert, intermediate, and expert, are shown. (**C**) The average arm speed across the reach cycle did not significantly differ among the three groups (one-way ANOVA, *F*_2,33_ = 0.37, *p* = 0.69). (**D**-**G**) The measurement and quantification of transverse muscle motions orthogonal to the direction of muscle fibers, captured by an ultrasound probe. (**D**) Sketches showing the placement of the ultrasound probe on the upper arm (left) and relative to a cross-section through the upper arm (right). The triceps, brachialis, and biceps muscle outlines are sketched in yellow, pink, and blue, respectively. The humerus bone is sketched in gray. The green dot represents the tracked point at the triceps-brachialis interface. The blue line represents the probe length (3.75 cm), relative to the cross-sectional sketch of an image from the visible human project[87]. (**E**) A representative ultrasound image, along with the motion path of the tracked point. (**F**) Speed profiles of the muscle interface for each participant group. (**G**) The average speed of the tracked point throughout the reach cycle was significantly different across the participant groups (one-way ANOVA, *F*_2,33_ = 5.42, *p* = 0.0092). Tukey’s post-hoc test showed that the experts’ muscle interface speed differed from that of the other groups (\*\**p* = 0.0084, *CI* = -1.68, -0.220 for expert vs non-expert; *p* = 0.060, *CI* = -1.36, 0.0231 for expert vs intermediate), but was similar between the non-expert and intermediate groups (p = 0.58, *CI* = - 0.406, 0.973). Here, ’n’ represents the number of participants, error bars and shaded areas represent S.E.M., and ’n.s.’ stands for not significant.

### Movement experts have reduced transverse muscle motions

We examined if experts’ muscles move less in the transverse plane orthogonal to the direction of muscle fibers. We focused on the interface between muscles, rather than the motions of points within muscles, as the latter include relative motions between muscle fascicles.

To measure transverse muscle motions, we used brightness mode (B-mode) ultrasound imaging to capture parts of the triceps and brachialis muscles in the upper arm (Fig. 2d,e, Fig. S3) and tracked a point at their interface toward the superficial end (Fig. 2d green circle). We then extracted the speed profiles of this triceps-brachialis interface during the reaching task (Fig. 2f) and observed a lower speed in the expert group than in the intermediate and non-expert groups, indicating reduced transverse motion in experts (Fig. 2g). The muscle interface speed was also more consistent in experts across repetitions of the reaching movement (Fig. S4a,b).

To better understand the motion of the muscle interface, we tracked additional points along the interface from the humerus bone to the fat layer. Superficial points closer to the skin had lower speeds (Fig. S4e,f), which was expected because tissue speed was measured relative to the skin. The muscle interface speed at the bone was similar across expertise levels (Fig. S4c,d). However, the muscle interface speed decreased faster in experts than in non-experts from the bone to the skin, and the speed profiles along the interface were similar between intermediates and non-experts (Fig. S4e,f).

#### Data interpretation

These data suggest that the control mechanisms in experts may better account for the constraint mapping between muscles and body segments, thereby resulting in slower (Fig. 2f,g, Fig. S4f) and therefore smaller, but more consistent (Fig. S4a,b) transverse muscle motions in experts compared to intermediates and non-experts.

### Movement experts have fewer physiological tremors

Physiological tremors were discovered and described well over a century ago [34] and they are considered to have no clinical significance [41,45]. We found that experts across disciplines have fewer tremors when they move (Fig. 3), establishing the significance of physiological tremors in relation to expertise.

**Fig. 3.**
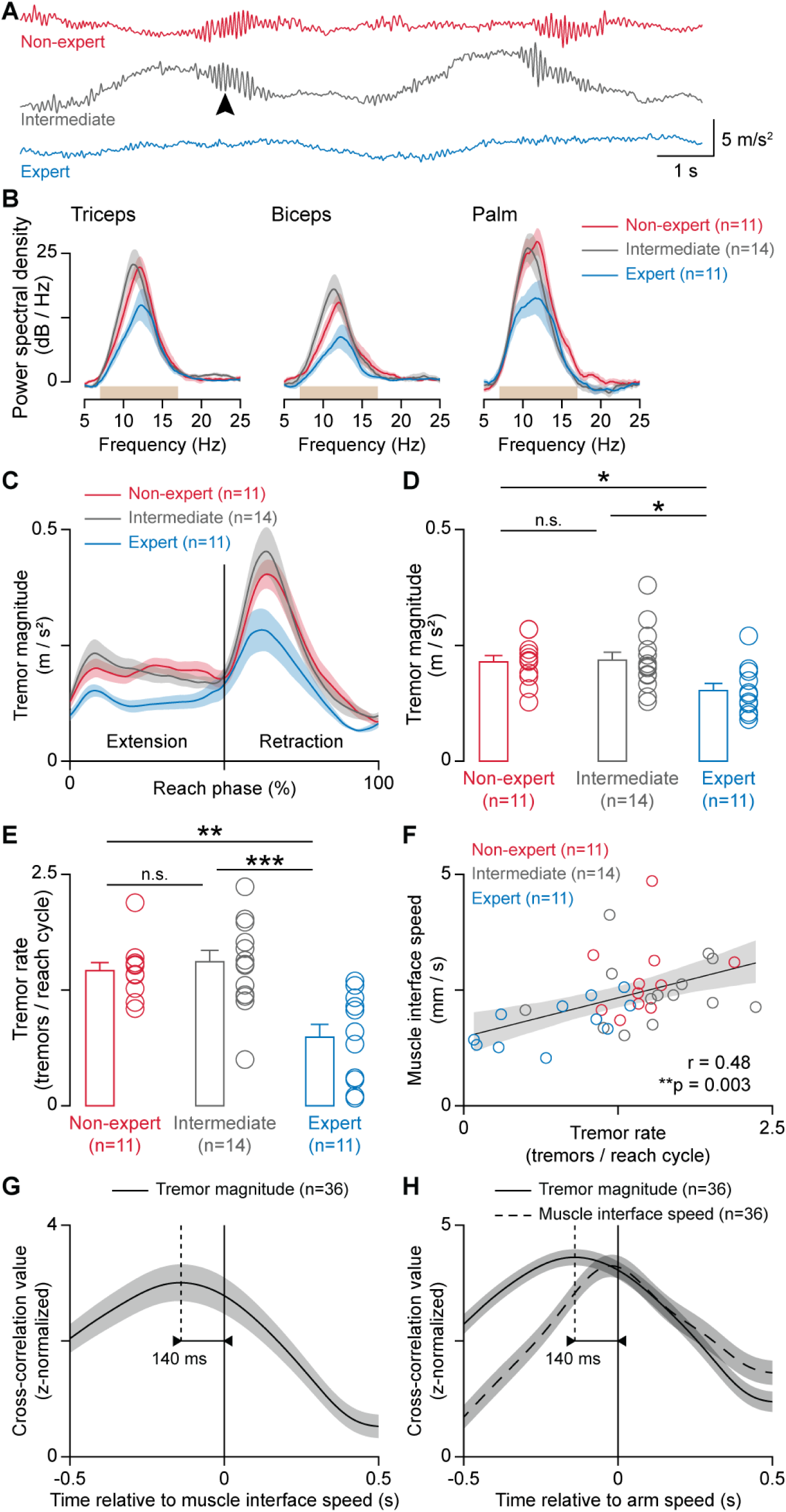
Movement experts have fewer physiological tremors. Accelerometer data recorded from the palm, triceps, and biceps of participants across different expertise levels (non-expert, intermediate, and expert) revealed distinct patterns. (**A**) Exemplary traces from each participant group (non-expert, intermediate, expert; top to bottom) captured by an accelerometer placed on the palm during the slow reaching task. The arrow points to a movement-induced tremor. (**B**) Power spectral density (PSD) analysis of accelerometer data from each of the three sensor locations, the triceps, the biceps, and the palm, for each subject group during the reaching task. The light brown boxes show the frequency band used to identify physiological tremors in the time domain (7-17 Hz). (**C**) Tremor magnitude, i.e., the amplitude of the accelerometer signal in the 7-17 Hz frequency band, as a function of the reach phase. (**D**) The average tremor magnitude across the reach cycle was significantly lower in the expert group compared to the other two groups (one-way ANOVA, *F*_2,33_ = 4.76, *p* = 0.015; Tukey’s post-hoc test, \**p* = 0.043, *CI* = 0.00174, 0.122 for non-expert vs expert; \**p* = 0.020, *CI* = 0.00898, 0.123 for intermediate vs expert), and was similar between non-expert and intermediate groups (*p* = 0.98, *CI* = -0.0607, 0.0529) (**E**) The tremor rate differed significantly across groups (one-way ANOVA, *F*_2,33_ = 12.3, *p* = 0.00010), with a lower tremor rate in the expert group (Tukey’s post-hoc test, \*\**p* = 0.0013, *CI* = 0.266, 1.17 for non-expert vs expert; \*\*\**p* = 0.00014, *CI* = 0.386, 1.24 for intermediate vs expert) and similar tremor rates in the non-expert and intermediate groups (*p* = 0.85, *CI* = -0.522, 0.331). (**F**) Correlation plot showing that participants with fewer tremors had a lower muscle interface speed during reaching (Pearson correlation, *r* = 0.48, \*\**p* = 0.0030). (**G**) Cross-correlation analysis showed that the strongest relationship between muscle interface speed and tremor magnitude occurred with a time offset of 140 ms, as indicated by the dotted vertical line. This finding showed that changes in tremor magnitude precede changes in muscle interface speed. (**H**) Similarly, the strongest relationship between arm speed and tremor magnitude occurred at the same offset of 140 ms (vertical dotted line), with changes in tremor magnitude preceding changes in arm speed (solid black trace). The peak correlation between arm speed and muscle interface speed (dashed trace) did not show a noticeable temporal offset. Here, ‘n’ represents the number of participants, error bars and shaded areas represent S.E.M., except in panel f, where the shaded area represents 95% confidence intervals, and ‘n.s.’ stands for not significant.

We used accelerometers to capture tremors across levels of expertise during the reaching task. Spectral analysis of accelerometer data (Fig. S5a-c) from the triceps, biceps, and palm showed increased power between 7 Hz and 17 Hz at every sensor location (Fig. S5d), with a distinct peak at approximately 12 Hz (Fig. 3a,b, Fig. S5c). Notably, the peak frequency did not differ across participant groups, but was lower at the palm (Fig. S5g-h). A time domain analysis showed that experts had lower signal amplitude in this frequency band during the reach cycle, compared to the non-experts and intermediates (Fig. 3c,d).

To discern whether experts had lower-amplitude tremors, fewer tremors, or both, we segmented the tremor events (Fig. S6). This analysis revealed that experts experienced tremors less frequently during reaching movements compared to non-experts and intermediates (Fig. 3e). The peak amplitude and average duration of the tremors were similar across all groups (Fig. S5e,f). Therefore, even though tremors occur less frequently in experts, when they do happen, the tremor properties are similar across non- experts, intermediates, and experts.

Reduced tremors and transverse muscle motions in experts are not explained by muscle size or by the size of the fat layer in the upper arm (Fig. S7). While participants who identified as female had a smaller muscle size (Fig. S7c) and larger fat layer size (Fig. S7g) than those who identified as male, there was no difference in these parameters across levels of expertise (Fig. S7b,f). There was no correlation between these tissue size parameters and muscle interface speed (Fig. S7d,h), or tremor rate (Fig. S7e,i).

*Tremors correlate with and precede transverse muscle motions.* Next, we investigated whether transverse muscle motions are related to tremors and found that individuals with lower muscle interface speed also had fewer tremors (Fig. 3f). We wondered whether transverse muscle motions play a role in triggering physiological tremors. Surprisingly, a cross-correlation analysis revealed that fluctuations in tremor magnitude preceded those in muscle interface speed by 140 ms (Fig. 3g). A similar analysis showed that tremor fluctuations also precede arm speed fluctuations, while the peak correlation between arm speed and muscle interface speed showed no noticeable temporal offset (Fig. 3h).

#### Interim discussion

Based on the data presented thus far, experts have reduced inefficiencies in muscle motions, both in terms of transverse muscle motions and tremors, compared to non-experts and intermediates. Interestingly, these inefficiencies are comparable between non-experts and intermediates, rather than decreasing with increasing expertise. This suggests multiple stages of expertise development from the perspective of muscle motions, with different aspects of muscle motions being optimized at each stage.

### In intermediates and experts, arm motions similar to those of non-experts are accompanied by smaller muscle length changes

We measured muscle length indirectly from ultrasound videos that captured the muscle’s cross-section. Since the volume of a muscle is constant [46–49], changes in muscle area reflect muscle length changes [50]. We measured the area of a portion of the triceps muscle over time by tracking four points within the triceps, and extracting the enclosed area (Fig. S8a,b). The rate of area change, and therefore the speed of muscle length change, was similar across participant groups (Fig. S8c-f), which we expected as a consequence of similar arm movement speeds across participant groups.

We then examined whether expertise influences the relationship between muscle length changes and arm movements. To analyze this relationship, we used a geometric formulation. Consider the path of a point that jointly represents the arm position and muscle length (Fig 8a). The angle of the path tangent quantifies the relative proportion of these movements. A larger angle indicates that <u>smaller muscle length changes</u> <u>correspond to arm motions of a given size, signifying efficacy</u>. This analysis showed significant differences between participant groups (Fig. 4c), especially near the transition between the extension and retraction phases (Fig. 4b). Therefore, participants in both the intermediate and expert groups demonstrated more effective translation of muscle length changes to arm movements during reaching, with smaller muscle length changes corresponding to arm movements of a given size.

**Fig. 4.**
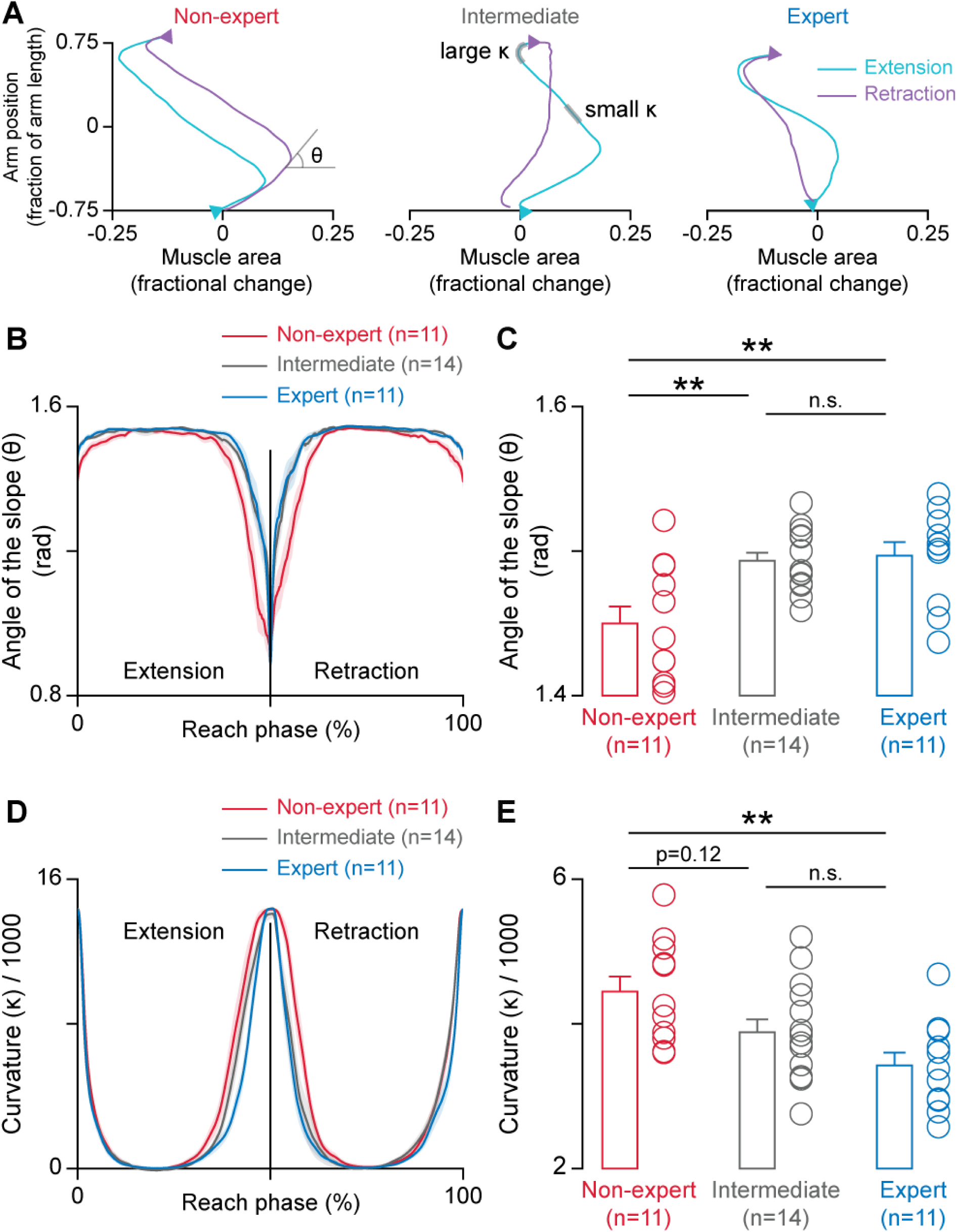
In intermediates and experts, smaller muscle length changes correspond to arm movements of a given size. (**A-C**) Proportion of muscle length change and arm movement during reaching. (**A**) Paths of normalized muscle area (proxy for muscle length) and normalized arm position from a representative participant in each of the non-expert, intermediate, and expert groups. The angle between the horizontal and the path’s tangent, θ, signifies the relative proportion of arm movements and muscle length changes. The curvature, κ, signifies the complexity of the relationship between arm movements and muscle length changes. Two segments of the path, one with a small κ and one with a large κ are highlighted in gray. (**B**) θ values as a function of the reach phase for each of the participant groups. (**C**) Average θ over the reach cycle varied significantly across the participant groups (one-way ANOVA, *F*_2,33_ = 7.75, *p* = 0.0017). Tukey’s post-hoc test showed that θ was significantly lower in non-experts (\*\**p* = 0.0048, *CI* = -0.0746, -0.0121 for non-expert vs intermediate, and \*\**p* = 0.0040, *CI* = -0.0799, -0.0138 for non-expert vs expert). θ values of experts and intermediates were similar (*p* = 0.96, *CI* = -0.0277, 0.0347). (**D**) Curvature (κ) values as a function of the reach phase for each of the participant groups. (**E**) Average κ values over the reach cycle varied significantly across participant groups (one-way ANOVA, *F*_2,33_ = 6.19, *p* = 0.0052). Tukey’s post-hoc test showed that the κ of non-experts was significantly greater than that of experts (\*\**p* = 0.0036, *CI* = 307.5, 1735) and trended to be greater than that of intermediates (*p* = 0.12, *CI* = -110.9, 1237). The κ of intermediates and experts was similar (*p* = 0.23, *CI* = -216.4, 1132). Here, ’n’ represents the number of participants, error bars and shaded areas represent S.E.M., and ’n.s.’ stands for not significant.

The path curvature quantifies an aspect of the interaction complexity between muscle length and arm movements (Fig. 4a). A higher path curvature indicates more frequent changes in the relationship between muscle length and arm movements during the reach cycle, and thus a more complex relationship. The average path curvature was higher for non-experts compared to intermediates and experts (Fig. 4d,e), indicating that there was a more complex relationship between muscle length changes and arm movements in non-experts.

#### Data interpretation

The concept of muscle tone can help interpret these results. In this context, we define a *toned or tensile muscle state* as a condition where even minimal muscle-generated force can cause slight movement of the joint. In simple terms, a toned muscle behaves like a taut rubber band, while a muscle that lacks tone resembles a slack rubber band. If a muscle lacks tone at rest, and tone is required during movement, we expect an initial shortening to produce tone before the muscle contributes to the motion of body segments. We might also expect this muscle to lose tone towards the end of the movement if tone is not needed to maintain the final position. This may explain why we observed less effective muscle length changes at the start and end of the reach phases in non-experts and why there was a more complex relationship between muscle length changes and arm movements in non-experts.

#### Interim discussion

As we observed, in intermediates, the effectiveness of muscle- length changes resembled that of experts, but the precision (in terms of transverse muscle motions and tremors) was similar to that of non-experts. If precision in muscle motions is learned, then these data suggest that the effectiveness of muscle length changes improves first, and subsequently, the precision of muscle motions increases as one progresses to an expert level. Alternatively, if precision is innate, then some individuals are predisposed to become experts. Therefore, we sought to understand whether tremors and transverse muscle motions can be voluntarily regulated. The results from this biofeedback experiment refute the possibility that precision is entirely innate.

### Tremor-based biofeedback reduces inefficient muscle motions

A subset of intermediate and expert participants performed a biofeedback task. Each participant viewed the accelerometer signal from the biceps, triceps, and palm sensors, one at a time, in real time while performing a reaching task (Fig. 5a). We asked them to minimize tremors as much as possible while maintaining a pace similar to that of the visual metronome from earlier trials.

**Fig. 5.**
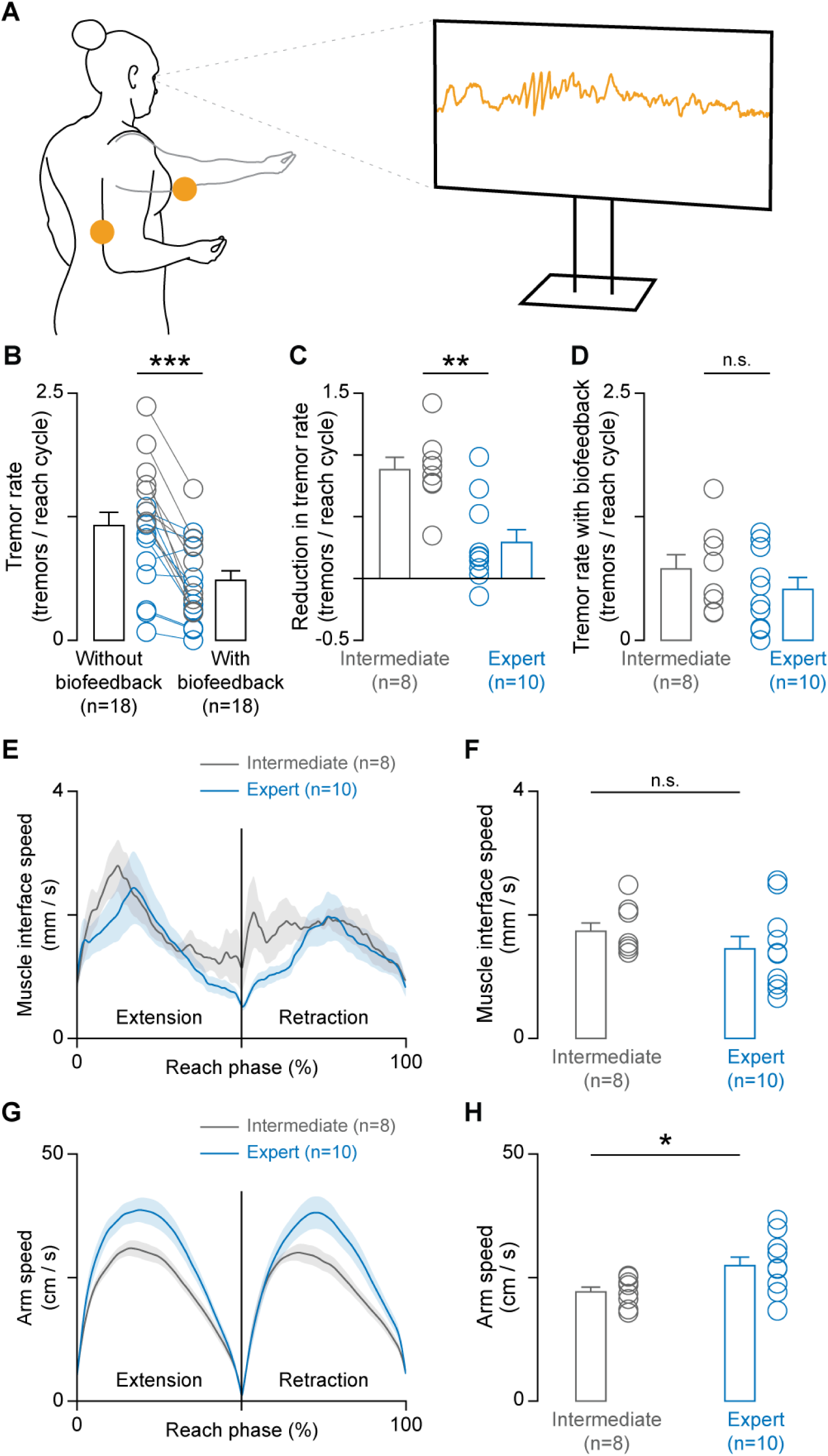
Biofeedback reduces tremors and muscle interface speed in intermediates, but comes at the cost of slower arm movements. (**A**) Experimental setup. Participants performed the reaching task while viewing accelerometer data on a screen in front of them. They were asked to minimize tremors and maintain a speed similar to that of the reaching task to the best of their ability. The accelerometer location on the triceps muscle is marked with yellow circles on the left, and a representative trace is depicted on the screen. (**B**) Tremor rate was significantly lower during biofeedback than during reaching with the visual metronome (paired *t*-test, *t*_17_ = 5.37, \*\*\**p* = 5.1 x 10^-5^), demonstrating the success of the biofeedback manipulation. The data corresponding to the intermediates (n=8) are shown in gray, and the data corresponding to the experts (n=10) are shown in blue. (**C**) During biofeedback, the tremor rate decreased more in intermediates compared to experts (*t*_16_ = 3.80, \*\**p* = 0.0016). (**D**) Tremor rate during biofeedback was similar between intermediates and experts (*t*_16_ = 1.043, *p* = 0.312). (**E**) Muscle interface speed as a function of the reach phase. (**F**) The average muscle interface speed did not differ between the intermediate and expert groups (*t*_16_ = 1.041, *p* = 0.313). (**G**) Arm speed as a function of the reach phase. (**H**) During biofeedback, the average arm movement speed was lower in the intermediate group than in the expert group (*t*_16_ = -2.36, \**p* = 0.032), suggesting that in order to reduce muscle interface speed and tremor rate to levels similar to those of experts, participants in the intermediate group slowed their arm movements. Here, ’n’ represents the number of participants, error bars and shaded areas S.E.M., and ’n.s.’ stands for not significant.

We observed that biofeedback successfully reduced the tremor rate (Fig. 5b). Participants in the intermediate group were able to reduce their tremor rate (Fig. 5c,d) and, consequently, muscle interface speed (Fig. 5e,f) to similar levels as experts.

However, this came at the cost of moving slower than experts (Fig. 5g,h).

### Data interpretation

These findings suggest a specific approach to achieving movement expertise. Practice should start slow enough to minimize inefficiencies (tremors and transverse muscle motions), and the movement speed should be gradually increased without introducing these inefficiencies.

## Discussion

Our key finding reveals that world-class experts across disciplines exhibit fewer inefficiencies in elastic tissue motions, thus establishing a connection between elastic mechanisms and general motor skill. Elastic mechanisms are typically studied for their role in efficiency and injury prevention [20,51–55], and this study broadens our understanding of elastic mechanisms.

This work challenges traditional views of athletic expertise. Current thinking in athletic expertise is that motor skills are highly specific, shaped by the unique demands and training methods of each discipline [11]. However, we have demonstrated that diverse athletic disciplines converge to affect fundamental aspects of movement: precision of elastic tissue motions and tremors. By doing so, we challenge the current thinking in athletic expertise: there are, in fact, general aspects to motor skill.

The definition of *general* plays a critical role in contextualizing these findings. We define "generality" of motor skill along two dimensions: athlete’s discipline (e.g., tennis/ballet) and movement type (e.g., specialized movements such as tennis serve and arabesque vs. everyday movements like reaching). Previous work has identified movement characteristics separating experts from non-experts, such as expert golfers exhibiting smoother golf swings [12]. Therefore, smoothness is a characteristic specific to both discipline (golf) and movement (golf swing). Some traits, like reduced motor variability, generalize across a wide range of disciplines but remain limited to specialized movements [56], such as expert golfers having less variability during golf swings, and expert tennis players having less variability during a forehand. It is particularly noteworthy that neither smoothness nor kinematic variability set apart experts across disciplines when performing everyday reaching movements (Fig. S2). In contrast, elastic tissue motions generalize beyond specialized movements across disciplines (Fig. 2,3,4), and in this sense, are indicators of general motor skill.

Our findings reveal that movement expertise profoundly influences how we move, even during everyday movements. Previously, we might have expected expert volleyball players to excel at jumps, gymnasts at balance. However, our research demonstrates that expertise has broader implications, influencing the quality of everyday movements. This is surprising as it is not immediately clear that perfecting an arabesque or serve- return should change how one reaches for a glass of water. It is even less obvious that the elastic tissue motions of experts in different disciplines – whether a ballet dancer or a volleyball player – would differ from non-experts *in the same way*. Yet, we found that experts across fields share common characteristics: fewer tremors, reduced transverse muscle motions, and more effective muscle-length changes.

The central ideology in approaching human movement – *muscles actuate bones to create movements* – places emphasis on the actuating function of muscles and guides the search for general indicators of skill towards cognitive and neural domains perception [57–62]. However, by showing that skill manifests in muscle motions not directly related to actuation, we show that general skill-related information is present in the body. Specifically, <u>skill is not simply related to the forces exerted by muscles to create joint</u> <u>movements, but instead it is related to the precision of muscle movements</u>.

Our findings demonstrate the importance of studying the motions of tissues within body segments, in addition to studying movement by capturing the motions of body segments. While previous studies have established that domain-specific skill information is evident in the more apparent movements of joints and body segments [12–15], we show that more general indicators of motor skill are evident in the less apparent motions of elastic tissues.

Certain indicators of general skill reported here (Fig. 2,3,4), such as fewer tremors, could signal superior *<u>quality of movement</u>*. We demonstrate that these indicators can be improved without rigorous training (Fig. 5), and this could have implications in improving movement quality and quality of life in the general population by reducing musculoskeletal injury risk and preserving movement quality in the aging population to reduce fall risk.

Our work is seemingly at odds with past work on generalized motor abilities [2,3]. Studies on general motor ability assumed that superior generalized motor ability would automatically lead to better performance across all motor tasks. However, despite the existence of famous multi-sport athletes, substantial evidence indicates that motor performance does not correlate across tasks, arguing against the existence of such a generalized motor ability [4–9]. Our research takes a different approach – by shifting the focus away from performance, we discover indicators of general motor skill.

Our perspective aligns with previous findings on general motor ability–we acknowledge that improving the precision of elastic tissue motions is perhaps insufficient to perfect an arabesque. While we have shown that voluntarily regulation of elastic tissue inefficiencies is possible (Fig. 5), the subsequent impact on performing specialized movements is unclear. Consider this analogy: improving movement quality to reduce the risk of repetitive stress injury does not make someone a better programmer. Clearly, domain-specific knowledge and practice remain essential to enhance performance.

Our data encourages us to re-consider the idea of motor educability, which is the ability to acquire motor skills [63–65]. To extend our programming analogy: It would be easier to improve programming skills when unencumbered by a repetitive stress injury. Similarly, it may be easier to acquire motor skills with superior indicators of general motor skill, such as fewer inefficiencies in elastic tissue motions.

The field of optimal motor control deals with formulating and optimizing costs, such as energy minimization that either predict or describe movement [18–22,66–68]. The field of sports expertise [11,69,70] has largely evolved separately from optimal motor control, despite an early appeal to cross-pollinate [71,72]. Our findings bring these two fields together to elucidate the fundamental nature of skill by examining optimizations related to tremor minimization or transverse muscle motion minimization.

Our findings can help movement practitioners reduce inefficiencies in muscle motions. Tremors, linked to transverse muscle motions (Fig. 3), can serve as a biomarker for inefficiencies. Inexpensive and easy-to-deploy accelerometers can detect tremors, offering a more affordable alternative to the costly ultrasound devices required to measure transverse muscle motions. Thus, our findings could lead to the creation and use of tremor-based biofeedback devices that enrich movement practice.

Finally, we expect these results to spark investigations into general aspects of motor skill, including the identification of physical substrates and mechanisms that support these general aspects of skill. The holistic organization of connective tissues in the human body [73–75] could provide an anatomical substrate for general motor skill.

Overall, we think that there are fundamental aspects of the motor system that remain under-exploited by untrained individuals, and those who dedicate their lives to honing motor skills may implicitly discover these facets of movement through their own sport and art forms.

### Limitations

Muscle length changes in this study are inferred indirectly from muscle area changes. While this is reasonable because muscle volume remains constant, this method could be less precise than tracking fascicle length changes. However, fascicle length tracking varies considerably in methodological difficulty depending on the muscles being studied. This method, while indirect, is more practical for generalizing across muscles. During conventional ultrasound imaging, the anatomical section being imaged changes slightly during movement. This general limitation associated with ultrasound imaging applies to our measurements as well.

We examined experts across nine different movement disciplines. While this provides a solid foundation for demonstrating generality across domains, the vast array of movement disciplines calls for future studies with larger sample sizes across a broader spectrum. Such expanded research would shed light on the nuanced relationships between expertise and tremors, muscle length changes, and transverse muscle motions. For instance, does the precision of arm muscle motions differ between experts in disciplines that emphasize upper body movements (such as racquet sports, fencing, volleyball, and rowing), and those that focus on lower body movements (like football, cycling, and sepak takraw)? Similarly, this study examines reaching as a representative everyday movement and doesn’t examine the full range of daily activities. Future research could delve into more nuanced variations of how skill generalizes across various activities of daily living.

### Future directions

This study advances the current understanding of the general nature of motor skill. It lays the groundwork for a mechanistic and causal investigation into general aspects of motor skill. We anticipate future research to specify the anatomical substrates that experts utilize more effectively than untrained individuals, uncover the mechanisms underlying general aspects of motor skill, and investigate how these mechanisms reduce injury risk and enhance performance.

## Conclusion

Our findings establish that elastic mechanisms have a role in general motor skill. This discovery expands our understanding of elastic mechanisms beyond their known roles in efficiency and injury prevention. Moreover, it challenges traditional views of athletic expertise and the fundamental nature of motor skill.

## Materials and Methods

### Experimental design

#### Participants

Data were collected from 37 healthy adults. Data from one participant were excluded prior to analysis because the participant added an extra pronation-supination motion of the hand to the reaching task. Among the remaining 36 participants, aged 26.1±7.2 years, 15 identified as male, 19 as female, and 2 as non-binary. Participants were categorized into 11 experts, 11 non-experts, and 14 intermediate based on the criteria detailed below. All subjects provided written informed consent according to MIT’s Committee on the Use of Humans as Experimental Subjects (COUHES).

#### Expertise categorization

The cohort of movement experts consisted of individuals from diverse backgrounds, including athletes and performance artists. To be considered an expert, a participant had to have achieved notable recognition in their field, such as placing in international competitions like the Olympics or major national competitions like the NCAA national championship. Additionally, performance artists who had showcased their talents on a global platform, such as by performing on a Broadway show, were included due to the absence of competitions in their domain. All individuals in the expert category were required to be actively competing and/or practicing their movement discipline at the time of their participation.

In this study, the cohort of movement experts included seven dancers representing various genres such as Latin and ballroom dance, Hip-Hop, modern dance, ballet, Bharatnatyam, and theater jazz. In addition, there was one Broadway performance artist and three sports athletes in the expert cohort, including a squash player, a volleyball player, and a rower.

Non-experts were individuals who did not receive any formal training or engage in the serious pursuit of any movement discipline. Participants with some formal training, such as regional athletes who did not meet the expertise criteria were categorized as intermediate.

#### Task description

In this study, participants performed slow reaching movements, taking approximately 6 seconds per extension-retraction cycle. They paced their movements using a visual metronome, which was a dot oscillating vertically on a screen in front of them at 1/6 Hz.

Participants were instructed to start and end their reaching movements with their hand next to their hip. They were then told to reach forward in front of them, following the pace of the metronome. They did this either with their palm facing up, as if giving something, or with their palm facing down, as if touching something. Each participant performed about 10 ’give’ cycles and 10 ’touch’ cycles.

Participants performed the task while standing, using a randomly selected arm and disregarding dominant side preference. Among the participants, 32 identified as right-dominant, 4 as left-dominant. 21 participants (approximately 58%) performed the task using their dominant side.

In addition to the reaching task, a subset of participants (eighteen) also performed a biofeedback task, in which they visualized the raw accelerometer signal in real time using the Delsys Trigno Discover software (Delsys, Inc., Natick, MA, USA). An experimenter first taught participants to visually identify tremors, which took about 2 minutes. Then, participants performed three biofeedback trials, visualizing the accelerometer signal from the palm, triceps, and biceps in a randomized order. They were unaware of which sensor was providing biofeedback during any trial. Participants were instructed to perform the task at a similar cadence to the reaching task while minimizing tremors. No further instructions on how to achieve this were provided.

### Data collection

As the participants performed the reaching task, data were acquired from their arm using motion capture, ultrasound, and accelerometers.

For motion capture, 3D positions of infrared reflective markers were tracked at 240 Hz using 24 Optitrack Prime13 cameras, and the data were synchronized with the accelerometer and ultrasound systems using eSync2 (NaturalPoint, Inc., Corvallis, OR, USA). The markers were affixed to the participants’ skin with kinesiology tape (Hampton Adams). A total of five markers were used to track arm movements and they were placed in the shoulder (acromion), upper arm (deltoid insertion into the humerus), elbow (lateral epicondyle), forearm (midpoint between the distal end of the ulna and the lateral epicondyle of the elbow), and hand (distal portion of the capitate on the dorsal surface). Motion capture data were acquired, labeled, and exported to a comma separated values (CSV) file format using Motive software (NaturalPoint, Inc., Corvallis, OR, USA) version 3.

Ultrasound data were obtained using a Cephasonics Cicada Ultrasound system (Cephasonics Ultrasound Solutions, San Jose, California, USA) coupled with a 7.5 MHz 128-channel linear ultrasound probe that was 3.75 cm long. The probe was affixed to the upper arm using a custom 3D-printed attachment and self-adhesive wrap. B-mode images were captured at an average rate of approximately 60 frames per second. The raw data were uniformly resampled at 60 Hz using nearest frame resampling. The ultrasound imaging provided transverse views encompassing the surface of the humerus bone, as well as segments of the brachialis and triceps muscles.

Wireless sensors from Delsys (Delsys, Inc., Natick, MA, USA) were used to collect acceleration data. These sensors were placed on the biceps, triceps, and abductor pollicis brevis muscles. The data were sampled at 148 Hz from the 3-axis accelerometers. The Trigno Discover software was used for data acquisition, real-time visualization during the biofeedback trials, and for exporting all the captured data into the CSV file format.

### Data analysis

Data analysis was performed using popular python packages such as Matplotlib and SciPy, and custom Python scripts. Custom-written, publicly available python packages (https://pypi.org/project/datanest, https://pypi.org/project/pyfilemanager) were used for file and data management. The custom code used for signal processing, generating bar graphs, and the user interface for labeling ultrasound videos are available publicly (https://github.com/praneethnamburi/pn-utilities).

We estimated arm speed from the 3D position data of the five tracked markers. Position data were low-pass filtered at 10 Hz. Then, these data were treated as a 15-dimensional vector at each sampled frame, and principal component analysis [76,77] was used to project the data along the axis with the most variance. The sign of this first principal component (PC1) was adjusted such that movement (slope of PC1) in the same direction as the shoulder-hand vector (extension) was positive, and movement in the opposite direction was negative (retraction). Arm velocity was defined as the first derivative of PC1, and arm speed was defined as the absolute value of this velocity.

To identify the start and end of each reach cycle, as well as the transition between the extension and retraction phases, we used arm velocity threshold crossings (Fig. S1). A reach cycle consisted of one extension followed by one retraction. The extension started when the arm velocity crossed 5 cm/s with a positive slope, and retraction started when it crossed -5 cm/s with a positive slope. The transition point was the midpoint of the arm velocity crossings with a negative slope, that is, the mid-point of when the arm velocity reduced below 5 cm/s and when the arm velocity fell below -5 cm/s. For reach-phase profiles (e.g., Fig. 2b,f, Fig. S2a, Fig. S4a,c), signals were resampled using linear interpolation to obtain 500 samples each for extension and retraction, collapsed over reach cycles using average (e.g., Fig. 2b,f) or standard deviation (Fig. S2a) for one participant, and then averaged across participants.

Movement smoothness (Fig. S2c-e) was quantified using the log dimensionless jerk [78]. This metric was calculated by taking the sum of the squared jerk (third derivative of position), multiplying it by the ratio of the cube of the reach cycle duration to the squared peak velocity, and taking the natural logarithm of the resulting product.

To quantify muscle length changes and transverse muscle motions, we tracked specific points in B-mode ultrasound images (Fig. S3). Points were tracked using a combination of optical flow (Lucas-Kanade) [27] and deep learning algorithms (DeepLabCut versions 2.3.7-2.3.9) [28,29].

First, to create training data for the deep learning model, we used a custom graphical user interface (GUI) written in Python. The GUI allowed us to follow specific landmarks in the ultrasound video with temporal continuity, as opposed to precisely identifying the same landmark from ultrasound images randomly sampled throughout the video. This step was important for creating reliable labels for the training dataset. We labeled 20-30 frames during one reach cycle, and augmented the training data using Lucas-Kanade optical flow with sigmoid correction [79] between labeled points.

Second, a ResNet-50 [80] architecture was trained for 500,000 iterations, with model snapshots saved every 50,000 iterations. The snapshot with the best test error was used to analyze the ultrasound videos. Model results for each tracked point were then visually inspected, and the model was refined using additional labeled frames. Separate models were trained for each participant based on results from previous work [79].

Due to frame-to-frame noise in ResNet-50 model results when tracking points in ultrasound videos [79], we used optical flow for post-processing. We created half-second tracklets starting at each frame with Lucas-Kanade and sigmoid correction, then averaged all tracklets to produce tracking results where the derivative (velocity) is less noisy compared to the derivative of the output from the ResNet-50 model.

We inferred muscle length changes from the changes in area of the muscle. To quantify the changes in the areas of the triceps and brachialis muscles, we tracked four points in each muscle and computed the enclosed area (Fig. S8a-f). The area (A_t_) was expressed as a fractional change from the area at the start of extension (A_0_), (A_t_-A_0_)/A_0_. The speed of muscle length changes was quantified using the absolute value of the time derivative of this normalized area.

To quantify transverse muscle motions (Fig. 2d-g, Fig. S4), we tracked a point at the interface of the triceps and brachialis muscles closer to the superficial side of the muscle, and extracted its speed. Tissue speeds were low-pass filtered with a cutoff frequency of 5 Hz prior to further analysis. Note that all tissue positions are in the reference frame of the ultrasound probe placed on the skin.

To further compare the motion of the muscle interface between participant groups (Fig. S4f), we tracked additional points along the interface of the triceps and brachialis muscles from where the interface intersects the humerus bone to the other end of the interface near the fat layer. We tracked 4 points on average along the muscle interface from each participant. To understand the speed of a tracked point as a function of its location on the muscle interface and the expertise level of the participant, we performed an ordinary least squares regression using the statsmodels python package [81]. The dependent variable was speed, and the independent variables included position along the muscle interface scaled between 0 (at the humerus bone) and 1 (superficial end near the fat layer), variables used to represent participant category (non-expert, intermediate, or expert), and variables used to represent interactions between position and participant group. The statsmodels package estimated the model coefficients as well as their significance. A statistically significant coefficient corresponding to an interaction term indicated whether the slope in one participant group was different from that in another group.

For physiological tremor-related examination of the accelerometer data, we used frequency domain and time domain analyses. Frequency domain analysis was used to determine the tremor frequency bandwidth and peak frequency (Fig. S5). The power spectral density was calculated using the Welch method [82], smoothed with a 1 Hz moving average filter, and averaged across participants for each sensor location. A third-order polynomial fit in log-log space was subtracted from the average power spectral density. The start and end frequencies of the tremor bandwidth were determined by threshold crossings at 20% of the peak power spectral density rounded to the nearest integer, resulting in a bandwidth of 7 to 17 Hz.

We used time domain analyses to extract tremor magnitude and identify tremor onset and offset times. The signal processing steps for extracting tremor magnitude from accelerometer signals were as follows: (1) band-pass filtering between 7 and 17 Hz using a finite impulse response filter [83], (2) detecting the envelope using the Hilbert transform [84], (3) computing the Euclidean norm of the envelopes from the three axes, (4) detrending using the adaptive iteratively reweighted Penalized Least Squares (airPLS) algorithm [85], and (5) smoothing using a 0.5 s moving average filter. Tremor power was extracted using the same steps, with an additional squaring step before smoothing.

Onsets and offsets of tremors were identified by threshold crossings on the derivative of tremor power. A single threshold value per sensor location was set across all participants using pilot data. When multiple onsets or offsets were detected consecutively, they were manually refined, and onsets / offsets that were detected without a corresponding offset / onset event were removed (Fig. S6).

We extracted the tremor rate, amplitude, and duration using the tremor onset and offset times. The tremor rate, measured as tremors per reach cycle, was calculated as the total number of tremors divided by the total number of reach cycles. The tremor amplitude was defined as the maximum tremor magnitude between onset and offset, and the tremor duration was the average duration from onset to offset.

To understand the temporal sequence of arm movements, transverse muscle motions, and tremors (Fig. 3g,h), we performed a cross-correlation analysis z-normalized using a permutation analysis. We performed the permutation analysis by computing a baseline distribution of cross-correlation values at each time lag. Given two signals, say tremor magnitude and muscle interface speed, across 20 reach cycles, we shuffled the reach cycles from one of the signals and computed the cross-correlation value. We repeated this procedure 1000 times, and used the mean and standard deviation of this distribution to z-normalize the true cross-correlation value between the two signals, that is, without any shuffling.

To explore the relationship between muscle length changes and arm movements (Fig. 4), we represented the normalized muscle area (A_t_) and the first principal component of the arm movement (PC1) geometrically (A_t_-PC1 path). We defined the efficacy of muscle length changes as the arctangent of the slope of the A_t_-PC1 path, represented as an angle (θ) between 0 and π/2. The complexity of the relationship between muscle length changes and arm movements was defined as the curvature (κ) of the A_t_-PC1 path.

### Statistics

Statistical tests were performed using the Scipy [86] and statsmodels [81] Python packages, as well as GraphPad Prism v10 (GraphPad Software, Inc.). Statsmodels was used to perform ordinary least squares regression and subsequent tests to determine the statistical significance of the model coefficients (Fig. S4f). GraphPad Prism was used to perform repeated measures analysis of variance (ANOVA; Fig. S5h).

Within-subject comparisons were made using paired *t*-tests. Group differences were detected using one-way ANOVA or repeated measures ANOVA, followed by Tukey’s HSD (honestly significant difference) post-hoc test. Two-sided *t*-tests were performed unless specified otherwise. Corrections for multiple comparisons were made using Bonferroni correction when appropriate. For all the results, the significance threshold was placed at α = 0.05 (\**p* <= 0.05, \*\**p* <= 0.01, \*\*\**p* <= 0.001). All the data are shown as the mean and standard error of the mean (S.E.M.) unless specified otherwise. The reported confidence intervals (CI) are for a confidence level of 95%. Python code and results sheets of statistical tests from GraphPad software detailing (wherever applicable) estimates of variance within each group, the effectiveness of pairing in the case of paired *t*-tests and repeated measures ANOVA, comparison of variances across groups, etc. are available upon request.

## Acknowledgments

This work was carried out in part through the use of MIT.nano Immersion Lab’s facilities. We thank Micha Feigin-Almon for configuring the ultrasound system. We thank Mariia Smyk, Andrea Leang, Kelly Wu, and Isabel Waitz for assistance with data collection and annotation. We thank Vincent Chen and Charles Williams for assistance with data annotation. We thank Jessica Rosendorf for feedback on data processing and writing. We thank Micaela Amaral for assistance with figure illustrations. We thank Zachary Mainen, Jeremy Wong, Dhruva Raman, and Edward Horng-An Nieh for comments on the manuscript structure. We thank all the participants for volunteering their time. Professional editing services were not used in the preparation of this manuscript. Large language models were used to make minor improvements to the writing, and were not used to generate any of the ideas presented in this work.

## Funding

Sekisui House (BA)

NCSOFT (BA)

MIT Portugal Program NORTE01-0247-FEDER-045910 (DF, HG, BA)

“la Caixa” Foundation (ID 100010434) fellowship LCF/BQ/EU22/11930097

(RPL) MIT Bose Fellows Program (PN, LD)

## Author contributions

Conceptualization: PN, AK

Investigation: PN, RPL, UMS, ER

Data curation: PN, RPL, DMF, UMS

Software: PN, RPL, DMF

Visualization: PN

Funding acquisition: PN, HG, BA, LD

Project administration: PN, BA

Supervision: PN

Writing – original draft: PN

Writing – review & editing: PN, RPL, DMF, LD

## Data and materials availability

The data that support the findings of this study are available from the corresponding author upon reasonable request. The custom code used for signal processing and data management, and the user interface for labeling ultrasound videos are available publicly (see Methods). All other code used for data processing are available from the corresponding author upon reasonable request.

## Supplementary Materials

**Fig. S1.**
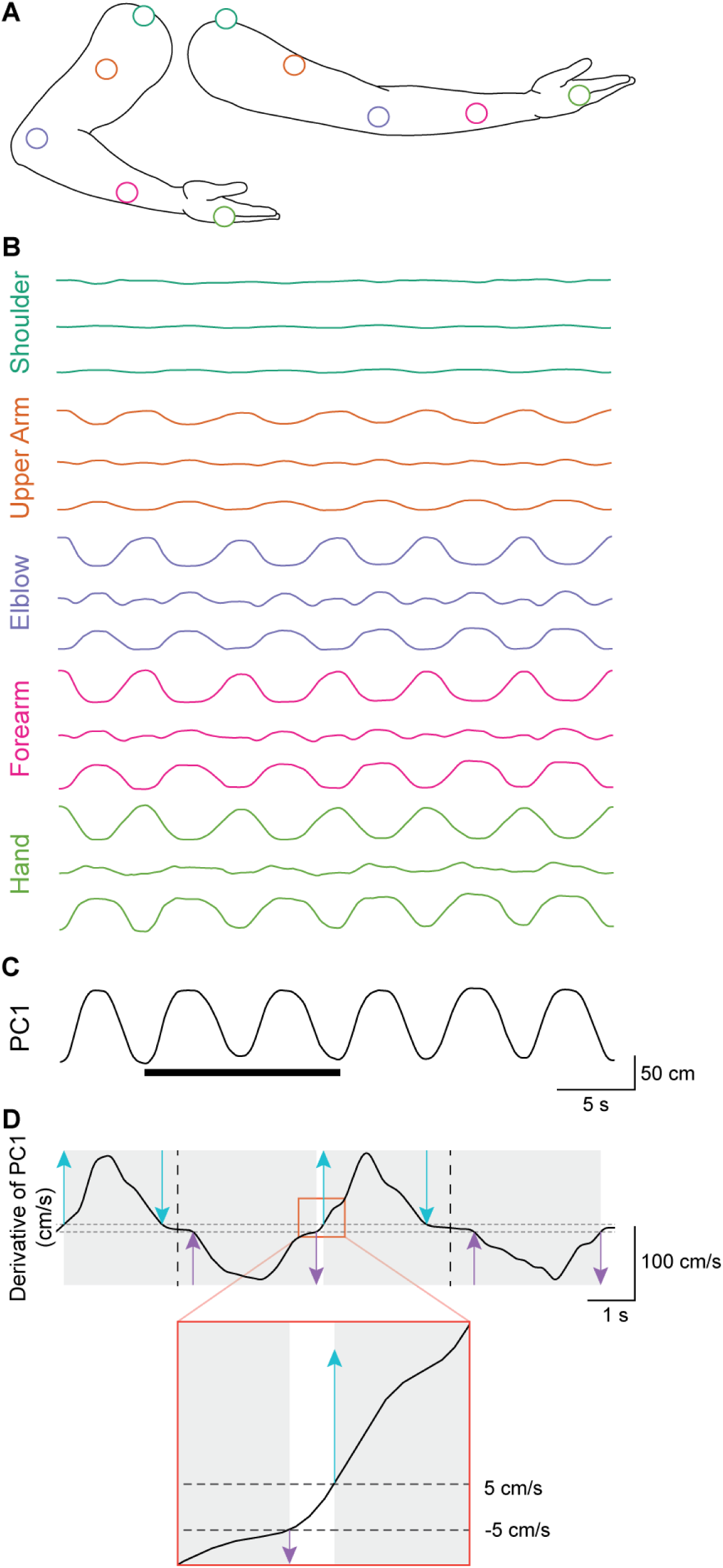
Reach cycle segmentation algorithm. (**A**) Locations of the five tracked points on the arm in the retracted (left) and extended (right) positions during the reach cycle. (**B**) Position data from the five tracked points over six reach cycles. Each tracked point has a set of three traces representing its position on each of the axes in 3D space with one trace for each of x, y, z orthogonal axes in Cartesian space. The top trace in each set shows position data in the anterior-posterior direction, the middle trace shows position data in the medial-lateral direction, and the bottom trace shows position data in the superior-inferior direction. (**C**) Position data are modeled as a 15-dimensional point, treating data from each time sample as one observation for Principal Component Analysis (PCA). The trace is the first principal component (PC1), which captures the reaching movement. The solid line below corresponds to the time span of the data shown in d. (**D**) The time derivative of the first principal component (solid trace) indicates arm speed. A 5 cm/s threshold (dashed horizontal lines) was used to detect the onset (up arrows) and offset (down arrows) of extension (cyan arrows) and retraction (lavender arrows). The transition between reach cycles (orange box) is magnified below to illustrate threshold-crossings. Each reach cycle spans from the onset of extension to the offset of retraction (gray boxes). The midpoint between the threshold-based offset of extension and threshold-based onset of retraction was considered the time point at which the movement switched from extension to retraction (dashed vertical lines).

**Fig. S2.**
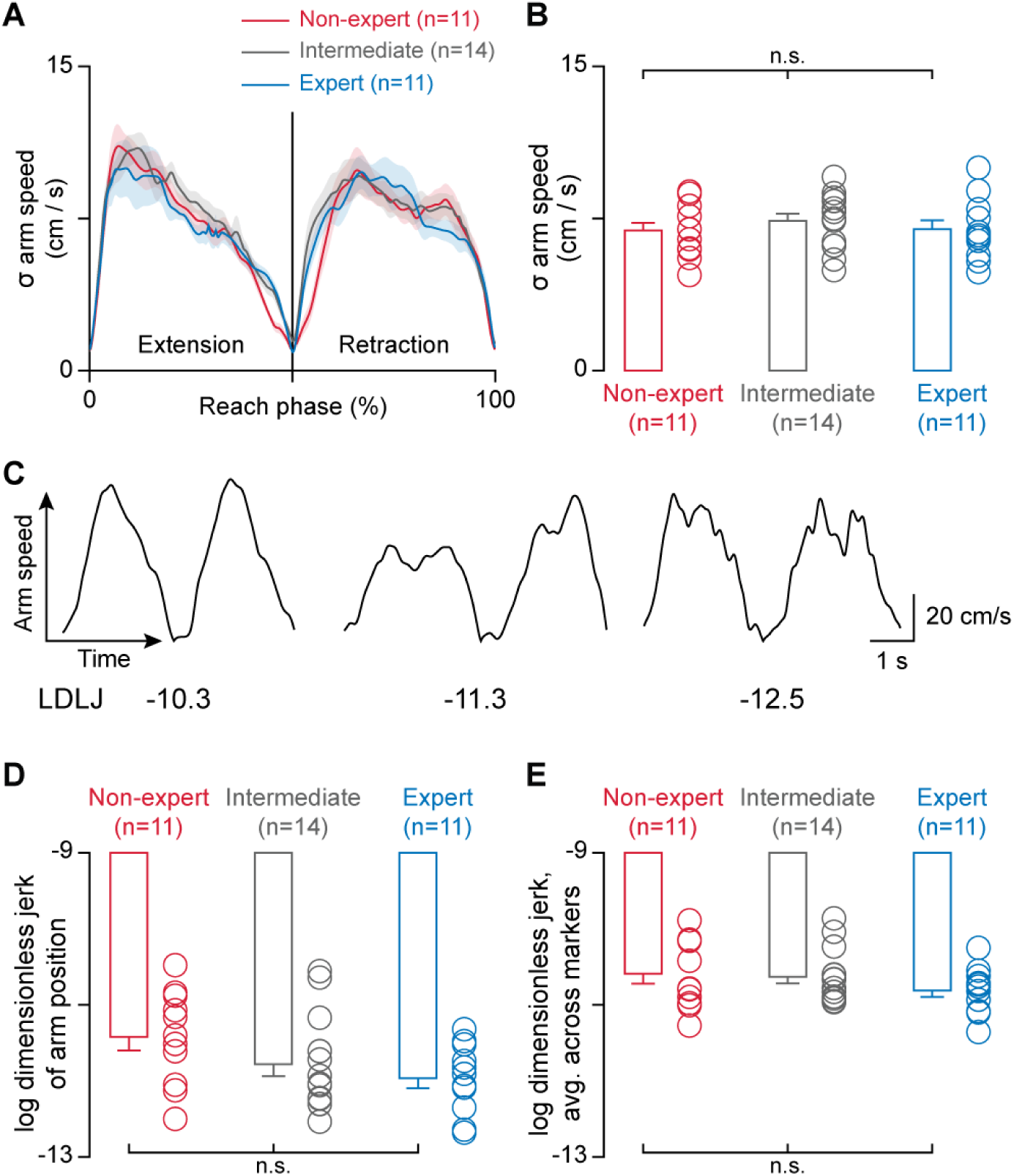
Arm movement metrics across levels of expertise. (**A**) Standard deviation (σ) of arm speed across trials as a function of the reach phase. (**B**) The variation in arm speed, measured using the standard deviation across reach cycles (σ), did not differ across participant groups (one-way ANOVA, *F*_2,33_ = 0.29, *p* = 0.75). (**C**) Three examples of arm-speed traces (top) from different participants, each showing one extension-retraction cycle, illustrating varying levels of movement smoothness. Movement smoothness was measured using log dimensionless jerk (LDLJ), with higher (less negative) values representing smoother movements (bottom). (**D**) There was no significant difference in the smoothness of arm movements (one-way ANOVA, *F*_2, 33_ = 2.59, *p* = 0.091). However, the trend suggests that expert movement may be slightly less smooth compared to that of non-experts during this reaching task. (**E**) The average movement smoothness across the tracked points on the arm also did not differ among the participant groups (one-way ANOVA, *F*_2,33_ = 1.12, *p* = 0.34). Here, ’n’ represents the number of participants, error bars and shaded areas represent S.E.M., and ’n.s.’ stands for not significant.

**Fig. S3.**
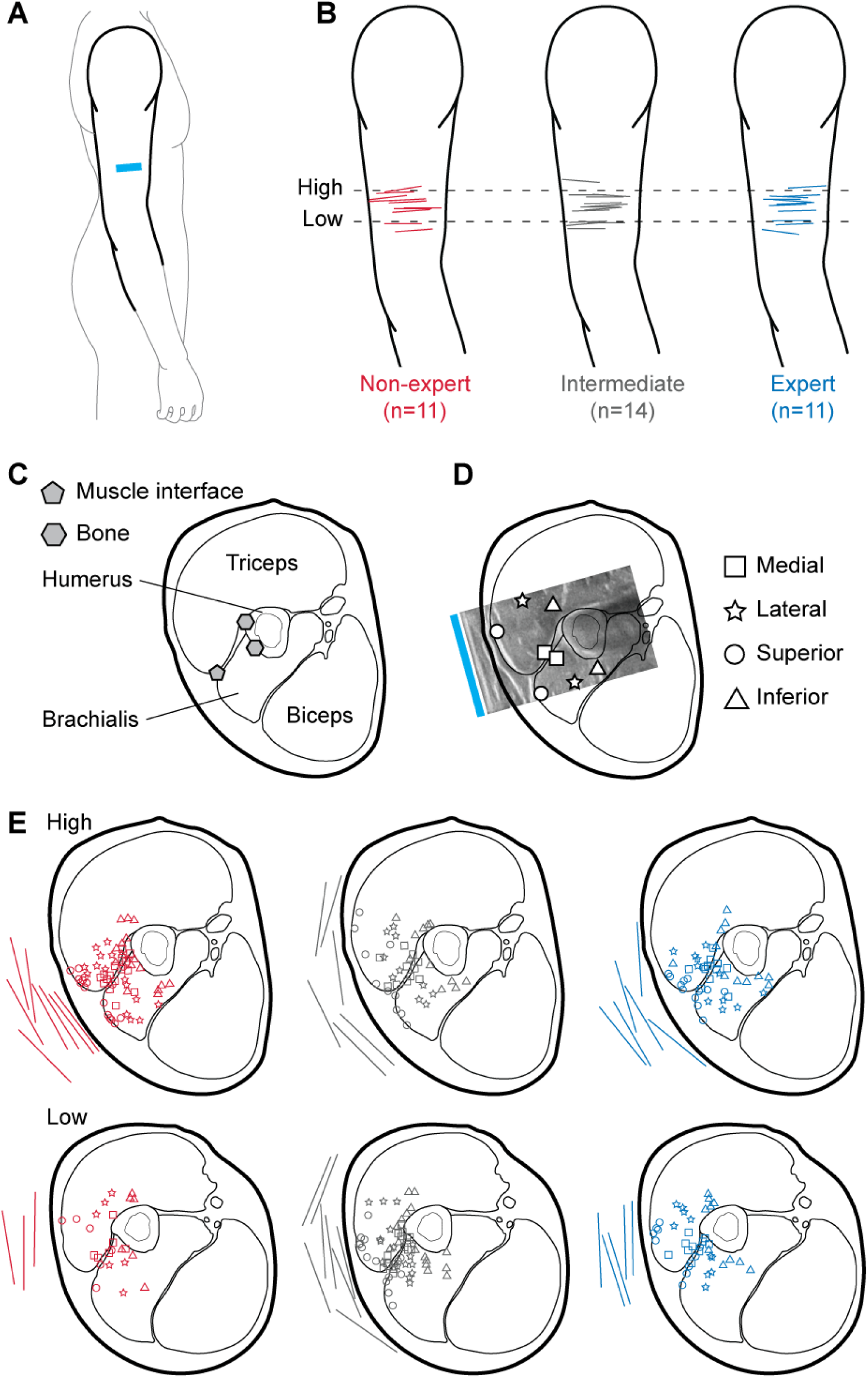
Ultrasound probe placements and locations of tracked points in the upper arm tissues. (**A**) A sketch showing the placement of the ultrasound probe (cyan bar) on the upper arm. (**B**) The exact placement of the probe for each participant was determined from a photograph of the probe on their arm. (**C-D**) Cross-sectional views of the upper arm illustrating the locations of tracked points. (**C**) A point at the interface of the brachialis and triceps muscles (gray pentagon) was tracked to measure transverse muscle motions. Additional points on the bone (gray hexagons) were tracked to infer the movements of the bone relative to the skin. (**D**) Four points were tracked in each of the triceps and brachialis muscles at the medial (square), lateral (star), superior (circle), and inferior (triangle) zones of the imaged muscle. The probe location relative to the cross-sectional diagram (cyan bar) was determined by fitting an ultrasound image taken at the start of the reach cycle to the cross-sectional diagram. (**E**) The probe placement on the circumference of the upper-arm cross section is depicted using lines, and the locations of the tracked points are shown using the symbols defined in d. The red, gray, and blue colors are for participants in the non-expert, intermediate, and expert groups, respectively. For convenience, data from participants with left arm probe placements are mirrored in this figure.

**Fig. S4.**
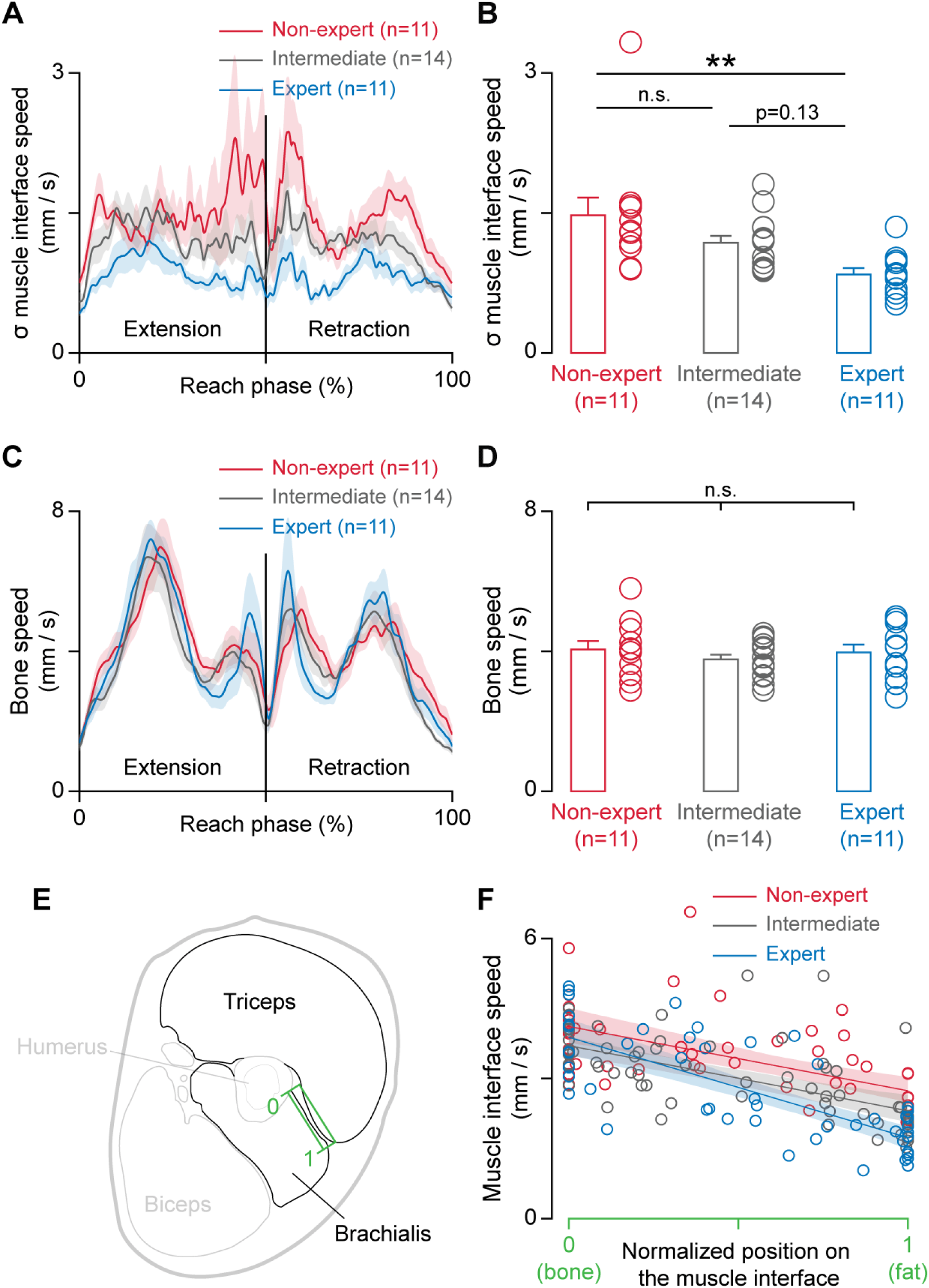
Movements of the interface between the triceps and brachialis muscles in the plane orthogonal to the muscle fibers. (**A**) Profiles of the variation in muscle interface speed, quantified as the standard deviation of muscle interface speed across reach cycles. (**B**) Experts had more consistent motions of the muscle interface across movement repetitions compared to non-experts and intermediates. The standard deviation of muscle interface speed was lower in movement experts (one-way ANOVA, *F*_2,33_ = 6.08, *p* = 0.0057; Tukey’s post-hoc test, \*\**p* = 0.0039, *CI* = -1.08, -0.188 for expert vs non-expert, *p* = 0.13, *CI* = -0.0822, 0.764 for intermediate vs expert, *p* = 0.22, *CI* = -0.128, 0.718 for non-expert vs intermediate). (**C**) The speed profiles of the point on the muscle interface that intersects with the bone. (**D**) The speed of the muscle interface at its intersection with the bone was similar across participant groups (one-way ANOVA, *F*_2,33_ = 0.53, *p* = 0.59), as opposed to the point closer to the fat layer (see Fig. 2f,g). (**E**) To better understand the behavior of the interface between the triceps and brachialis (green box), we sampled points not only at the two ends of the interface, that is, at the point intersecting with the bone (0) and the point close to the fat layer (1), but also in between these points along the muscle interface. (**F**) The speed of the points diverged from the bone towards the fat across the three groups. We tracked 49, 67, and 50 points from 11 non-experts, 14 intermediates, and 11 experts, respectively, along the muscle interface. We then performed an ordinary least squares regression with the speed of the point as a function of the position along the muscle interface, the participant category, and the interaction between the participant category and the muscle interface position. The interaction term for intermediates relative to non-experts was not significant (coefficient = -0.29, *p* = 0.31, *CI* = -0.861, 0.273), but the interaction term for experts relative to non-experts was significant (coefficient = -1.05, *p* = 0.0013, *CI* = -1.68, -0.416). This means that the slope of the regression line was different for experts compared to non-experts, and the slope of the regression line for intermediates was similar to that of non-experts, consistent with the findings reported in Fig.1h. Here, ’n’ represents the number of participants, error bars and shaded areas in panels a-d represent S.E.M., shaded areas in panel f represent 95% confidence intervals, and ’n.s.’ stands for not significant.

**Fig. S5.**
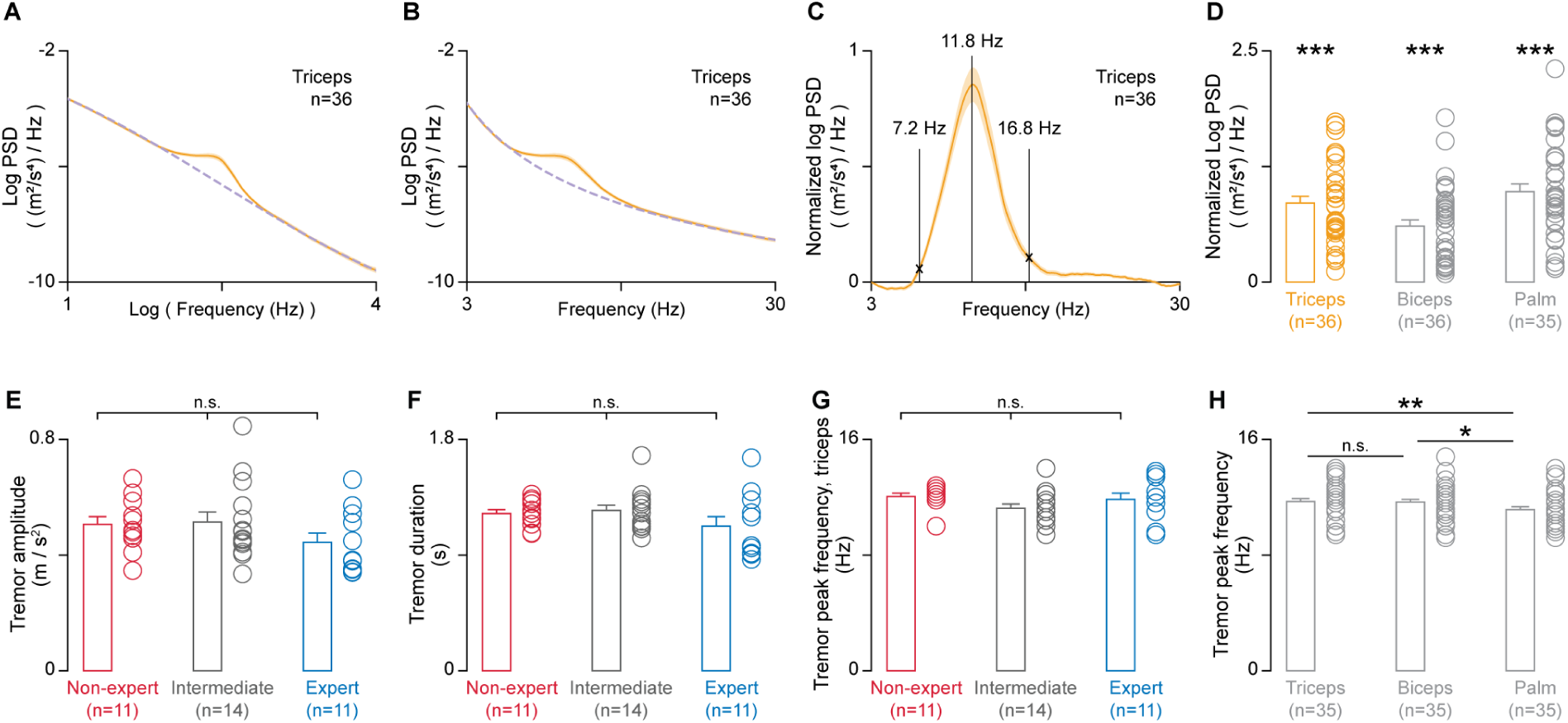
Spectral analysis of physiological tremors. (**A-D**) Tremor frequency band estimation. (**A**) The average power spectral density (PSD) from an accelerometer placed on the triceps (solid trace) in log-log space. A third-order polynomial was fit in log-log space (dashed trace). (**B**) Power spectral density and model fit visualized in linear-log space. (**C**) Normalized power spectral density, where the model fit is subtracted from the data, was used to estimate the frequency band (7.2 Hz to 16.8 Hz) and the peak frequency (11.8 Hz) at each sensor location. (**D**) At each sensor location, the power spectral density was significantly greater than the model fit (one-sample *t*-tests, *t*_35_ = 11.4, \*\*\**p* = 8.2 x 10^-13^ for the triceps, *t*_35_ = 8.93, \*\*\**p* = 4.5 x 10^-10^ for the biceps, *t*_34_ = 11.7, \*\*\**p* = 6.0 x 10^-13^ for the palm). Reported p-values were adjusted using Bonferroni correction for multiple comparisons. (**E-H**) Tremor properties. (**E**) Tremor amplitude, defined as the maximum tremor magnitude between tremor onset and offset, did not differ significantly across participant groups (one-way ANOVA, *F*_2,33_ = 1.26, *p* = 0.30). (**F**) Tremor duration did not differ significantly across participant groups (one-way ANOVA, *F*_2,33_ = 1.49, *p* = 0.24). (**G**) Peak frequency of the tremors at the triceps did not differ significantly across participant groups (one-way ANOVA, *F*_2,33_ = 1.75, *p* =0.19). The results were similar for the biceps and palm sensor locations (not shown; one-way ANOVA, *F*_2,33_ = 0.72, *p* = 0.50 for biceps; *F*_2,33_ = 1.26, *p* = 0.30 for palm). (**H**) The peak frequency of tremors was lower in the palm than in the triceps and the biceps (repeated measures ANOVA, *F*_2,34_ = 6.47, *p* = 0.0027; \*\**p* = 0.0052, *CI* = 0.147, 0.984 for triceps vs palm, \**p* = 0.011, *CI* = 0.101, 0.939 for biceps vs palm; *p* = 0.96, *CI* = -0.373, 0.464 for triceps vs biceps). Here, ’n’ represents the number of participants, error bars and shaded areas represent S.E.M., and ’n.s.’ stands for not significant.

**Fig. S6.**
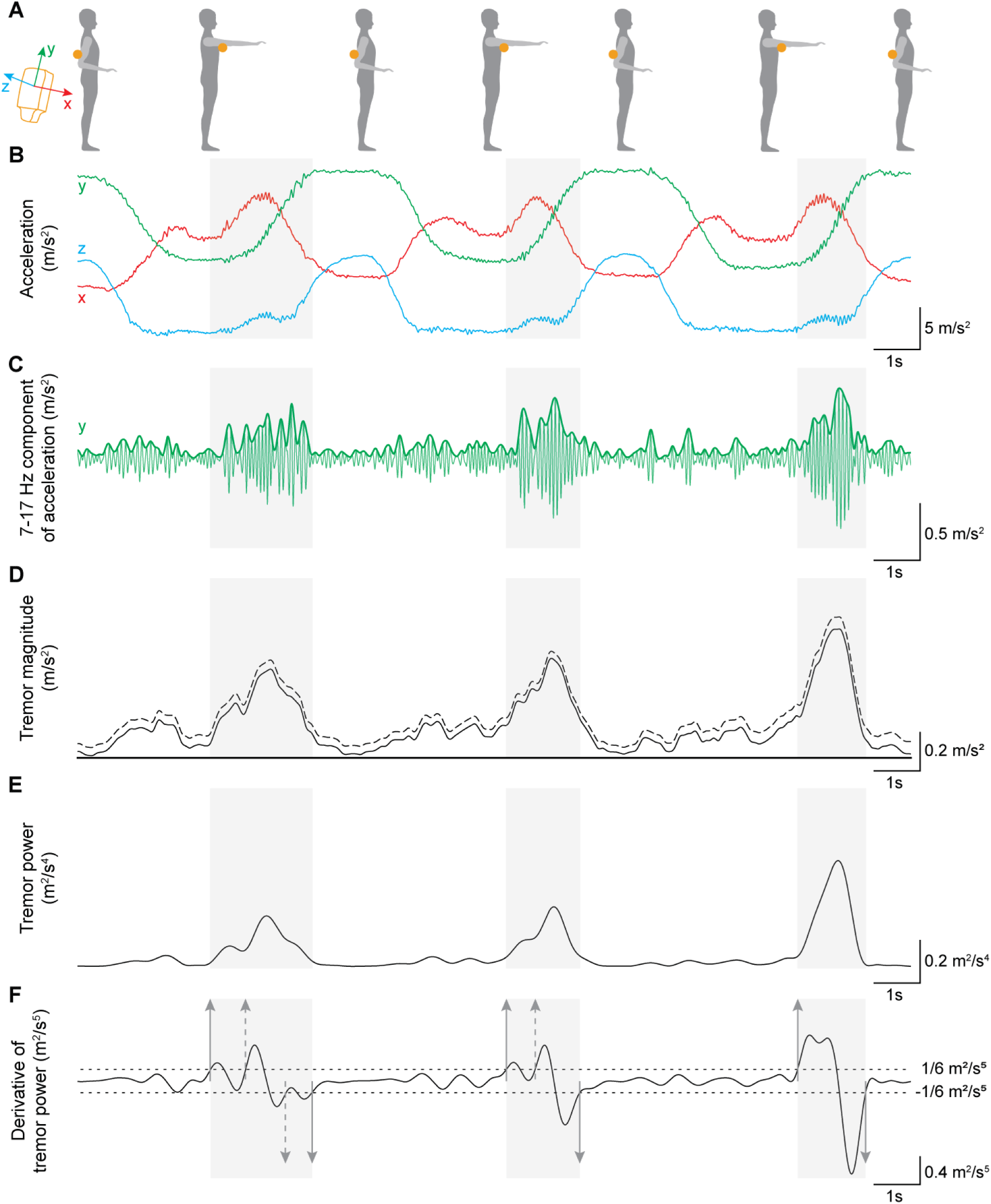
The tremor segmentation algorithm, which identifies tremor onsets and offsets, is based on threshold crossings on the derivative of tremor power. (**A**) An illustration of the reaching task. Yellow circles represent the placement of the 3-axis accelerometer, and the axis-convention of the accelerometer is shown on the left. (**B-F**) Signal processing steps involved in tremor segmentation. Light gray boxes represent identified tremors, which are detected after step (f), but are reproduced on all the panels for illustrative purposes. (**B**) Representative raw data traces recorded from the 3-axis accelerometer placed over the triceps muscle during the slow reaching task. The labels x, y, and z refer to the axes of the accelerometer. (**C**) Data from each axis were bandpass filtered between 7 and 17 Hz using a bandpass finite impulse response filter, and the envelope was detected. In this panel, the bandpass filtered signal from the y-axis is shown as a green trace, and the envelope is shown as a bold green trace. (**D**) Tremor magnitude was computed as the norm of the envelopes from each of the three axes (dashed trace), and the signal from the entire trial, which lasted approximately 2 minutes, was then de-trended using the adaptive iteratively reweighted Penalized Least Squares (airPLS) algorithm (*78*) (solid trace). (**E**) Tremor power, defined as tremor magnitude squared. (**F**) Thresholds (dashed horizontal lines) were imposed on the derivative of tremor power (solid trace) to identify onsets and offsets (gray arrows). In instances where two onsets were detected before an offset, one of the onsets was manually removed (dashed up arrows). Similarly, if two consecutive offsets were detected, one of the offsets was manually removed (dashed down arrow). A single threshold value was applied consistently across all participants for a given sensor location, determined based on pilot data.

**Fig. S7.**
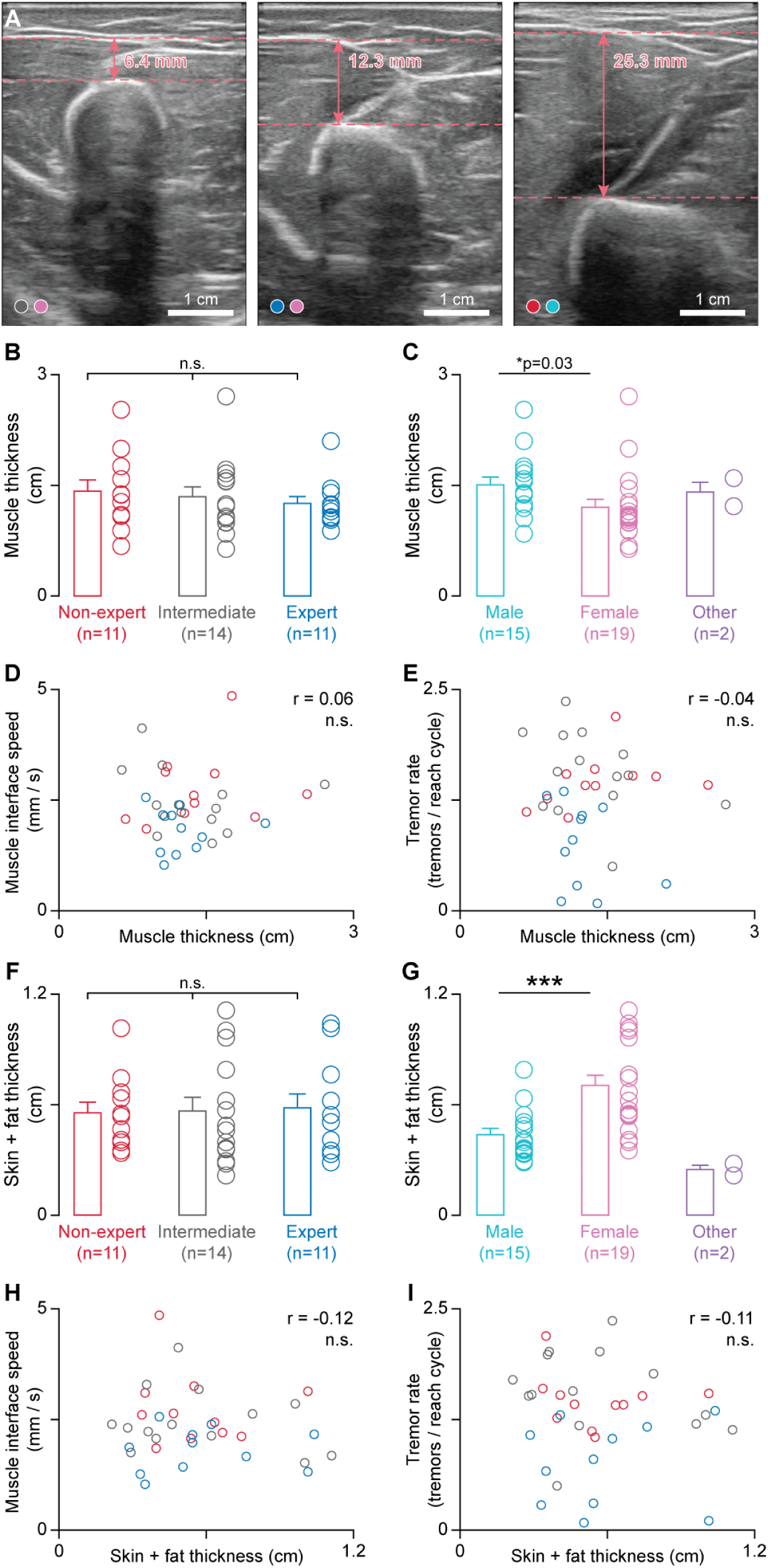
Muscle size and fat layer size do not correlate with transverse muscle motions or tremors. (**A**) Representative ultrasound images from three participants illustrating variation in muscle size. All three images were taken at the start of a reach cycle. The color of the first circle at the bottom left corner indicates the expertise level of the participant, and the second circle indicates gender. For color key, see panels B and C. Ultrasound images from left to right: intermediate female, expert female, and nonexpert male. Muscle thickness was measured as the distance from the top of the humerus bone to the bottom of the fat layer, with both landmarks visually identified, and represented using horizontal dashed lines. (**B**) There was no difference in muscle thickness across expertise levels (one-way ANOVA, *F*_2,33_ = 0.35, *p* = 0.71). (**C**) Muscle thickness of female participants was lower than that of male participants (one-sided *t*-test, *t*_32_ = 1.95, \**p* = 0.030). Since there were only two non-binary participants, we did not perform statistical analyses on this group. (**D**) Muscle thickness did not correlate with the speed of the point at the brachialis-triceps interface (Pearson correlation, *r* = 0.058, *p* = 0.74). (**E**) Muscle thickness did not correlate with tremor rate (*r* = -0.042, *p* = -0.81). (**F**) Similarly, the combined thickness of the skin and fat layers did not vary with expertise (one-way ANOVA, *F*_2,33_ = 0.033, *p* = 0.97). (**G**) Combined thickness of the skin and fat layers was different between male and female participants (two-sided *t*-test, *t*_32_ = -3.73, \*\*\**p* = 0.00074). (**H**) Combined thickness of the skin and fat layers did not correlate with the speed of the point at the brachialis-triceps interface (*r* = -0.12, *p* = 0.49). (**I**) Combined thickness of the skin and fat layers did not correlate with the tremor rate (*r* = -0.11, *p* = 0.52). Here, ’n’ represents the number of participants, error bars represent S.E.M., and ’n.s.’ stands for not significant.

**Fig. S8.**
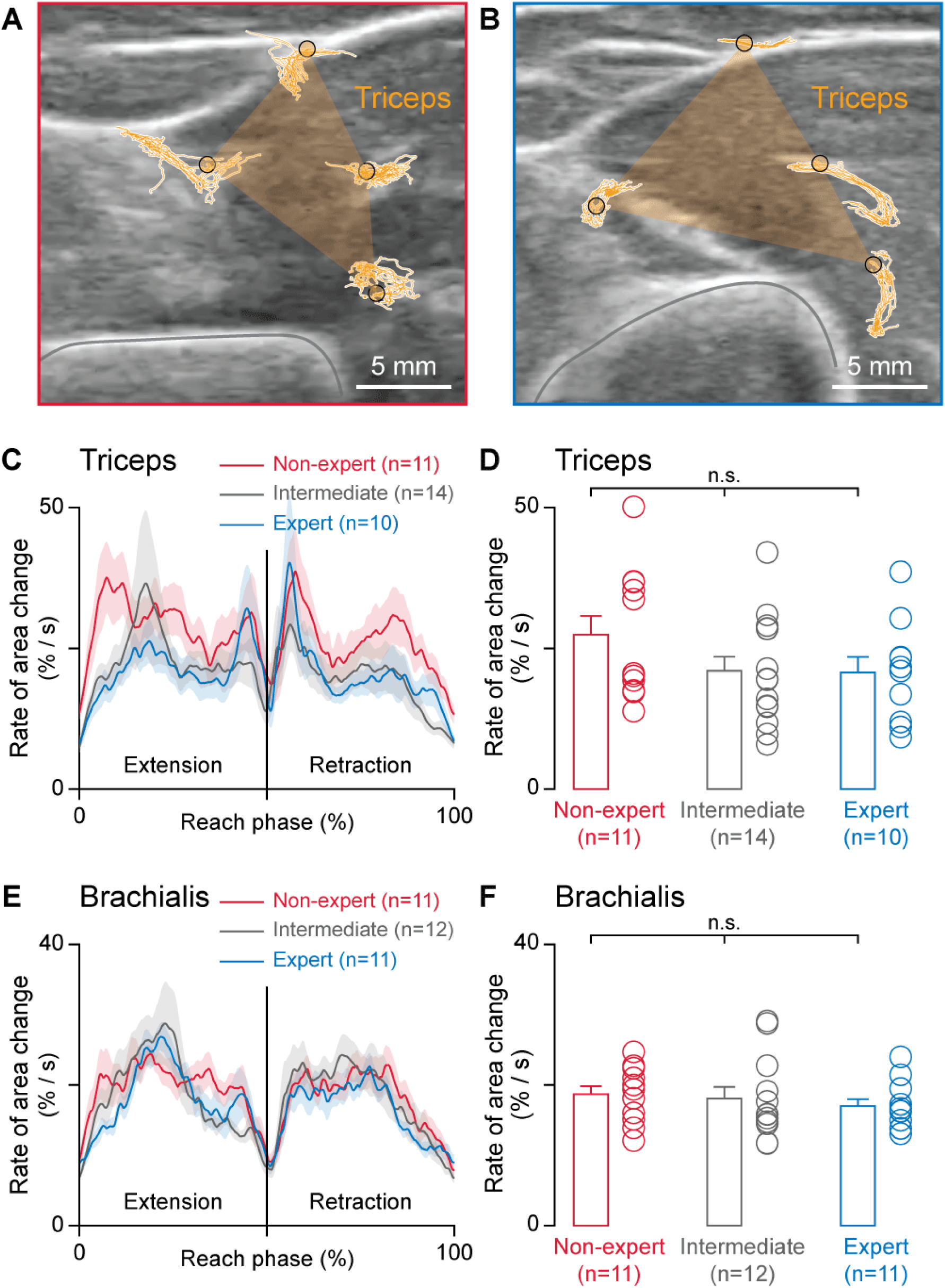
The speed of muscle length changes is similar across participant groups. (**A**) Representative tracking data in the triceps muscle of a non-expert. Overlaid on the B-mode ultrasound image are the four tracked points (yellow circles with black outline) and their tracked positions (yellow traces on white background). The area enclosed by the polygon formed by these points (yellow transparent overlay) is used to quantify muscle length changes. This area is normalized by the average area at the start of reach cycles so that the relative distances between the tracked points do not influence the results. The gray trace outlines the humerus bone. Note that the ultrasound image shown here is clipped to highlight the tracked points. (**B**) Representative tracking data in the triceps muscle of an expert. (**C**) The speed profiles of the triceps’ muscle length changes, quantified as the rate of change in area enclosed by the tracked points in the ultrasound video of the triceps. (**D**) The rate of change in the triceps area, averaged over the reach cycle did not differ across participant groups (one-way ANOVA, *F*_2,32_ = 1.55, *p* = 0.23). (**E**) The speed profiles of the brachialis’ muscle length changes. (**F**) The rate of change in the brachialis area, averaged over the reach cycle did not differ across participant groups (one-way ANOVA, *F*_2,31_ = 0.37, *p* = 0.69). Here, ’n’ represents the number of participants, error bars and shaded areas represent S.E.M., and ’n.s.’ stands for not significant.

**Movie S1. Contrasting muscle motions between a non-expert and an expert using representative examples.**

(**A**) The reaching task performed by a non-expert, with animated paths of markers placed on the hand, elbow, upper arm, and shoulder overlaid.

(**B**) An ultrasound video from the upper arm, capturing the interface between the triceps and brachialis muscles. Animated paths of three points along the interface: near the bone (green), in the middle (orange), and near the fat layer (lavender).

(**C**) Traces from a 3-axis accelerometer placed on the triceps showing small-amplitude oscillations. These are physiological tremors. Gray boxes represent physiological tremors detected through signal processing (Fig. S4). The solid black line in the middle marks the time corresponding to the task and ultrasound videos.

(**D-F**) Reaching task, ultrasound video, and accelerometer traces from a movement expert with slower muscle interface speed and fewer tremors.

